# Core passive and facultative mTOR-mediated mechanisms coordinate mammalian protein synthesis and decay

**DOI:** 10.1101/2024.07.04.602056

**Authors:** Michael Shoujie Sun, Benjamin Martin, Joanna Dembska, Cédric Deluz, David M. Suter

## Abstract

The maintenance of cellular homeostasis requires tight regulation of proteome concentration and composition. To achieve this, protein production and elimination must be robustly coordinated. However, the mechanistic basis of this coordination remains unclear. Here, we address this question using quantitative live cell imaging, computational modeling, transcriptomics, and proteomics approaches. We found that protein decay rates systematically adapt to global alterations of protein synthesis rates. This adaptation is driven by a core passive mechanism supplemented by facultative changes in mTOR signaling. Passive adaptation hinges on changes in the production rate of the machinery governing protein decay and allows partial maintenance of the cellular proteome. Sustained changes in mTOR signaling provide an additional layer of adaptation unique to naïve pluripotent stem cells, allowing near-perfect maintenance of proteome composition. Our work unravels the mechanisms protecting the integrity of mammalian proteomes upon variations in protein synthesis rates.

## Introduction

Proteins are the central macromolecular components of the cellular machinery. The cellular proteome must be constantly turned over to eliminate misfolded or damaged proteins, maintain proteome homeostasis, and respond to external stimuli. Protein turnover relies on the combined activities of protein synthesis and decay. The rate of protein synthesis is influenced by mRNA concentration and translation rate, whereas protein decay is affected by degradation and dilution through cell growth and division. The ubiquitin-proteasome system (UPS) and autophagy are responsible for the active degradation of proteins and protein complexes. The UPS handles the degradation of most cellular proteins, while autophagy primarily engages in protein degradation during cellular stress^1^. The rates at which proteins are synthesized and degraded can be adjusted to meet cellular needs, regulating the pace of proteome renewal^2,3^ for specific cellular function^4–6^ or to achieve a specific growth or division rate^2,7,8^. The contributions of degradation and dilution to protein-specific decay vary widely depending on their degradation rates; for example, short-lived proteins mostly decay through proteasomal degradation, while proteins with half-lives longer than the cell cycle mainly decay by dilution.

The synthesis of new proteins is a highly energy-consuming process that requires a continuous supply of amino acids^9,10^. In multicellular organisms, these resources are subject to large fluctuations and can affect protein turnover rates^11–14^. For example in humans, food intake can trigger an massive increases in protein synthesis rates (up to 100% in the case of muscles ^15^). Global protein synthesis rates also change as a function of developmental and differentiation stages^16,17^, in response to pathological stimuli such as viral infection^18^ or lipopolysaccharide^19^. The Integrated Stress Response (ISR) and the mechanistic/mammalian Target Of Rapamycin (mTOR) signaling pathways regulate global rates of protein synthesis and decay^20–23^. The ISR can modulate protein synthesis rates in response to cellular stresses, including alterations in protein degradation rates, through the segregation of mRNAs into stress granules to inhibit their translation^24,25^. The mTOR pathway stimulates protein synthesis through increased production of ribosomal proteins and by directly enhancing translation rates^26^. Its role in regulating protein decay is more controversial. While mTOR inhibition was shown to increase protein degradation rates acutely (within one hour)^27,28^, the opposite effect was reported by another study focusing on longer time scales^29^.

Protein synthesis and decay rates need to be tightly coordinated to maintain proteome concentration and composition within a narrow range, thereby ensuring the maintenance of cellular functions^30^. While rates of protein synthesis and decay were shown to be positively correlated in cultured cells^31^, the mechanisms underlying this coordination are unknown. Here, we show that changes in protein synthesis rates direct the adaptation of protein decay rates through a core passive and a facultative, mTOR-driven active mechanism, and we reveal their implications on the control of proteome concentration and stoichiometry.

## Results

### Protein decay rates adapt to changes in protein synthesis rates

To monitor protein synthesis and degradation rates in live cells, we used the Mammalian Cell-optimized Fluorescent Timer (MCFT)^31^, a translational fusion between the fast-maturing sfGFP and the slow-maturing mOrange2 fluorescent proteins (Fig.1A). Because of these differences in maturation rates, the relative changes in green (sfGFP) and red (mOrange2) fluorescence contain information about changes in synthesis and degradation rates^31–33^ (Supplementary Text). We fused the MCFT to a SNAP-tag, which allows the orthogonal measurement of degradation rates upon pulse-labeling with a SNAP-tag dye^31,34,35^. We additionally fused a PEST element to the C-terminus, resulting in a high degradation rate (*k_deg_*) (Fig.S1A) of the fusion protein (<u>S</u>hort-<u>L</u>ived <u>T</u>imer-<u>S</u>NAP or SLT). To infer changes in MCFT turnover from fluorescence trajectories, we used a mathematical model (STAR Methods, Supplementary Text) to compute the protein synthesis rate (*S*) and distinguish protein decay by active degradation (*k_deg_*) from decay through dilution occurring upon cell division (*k_dil_*) (Fig.S1B). From this model, an analytical relationship between the sfGFP to mOrange2 fluorescence ratio (called G/R hereafter) and the decay rate (*k = k_dil_ + k_deg_*) of the MCFT can be derived at steady state.

**Figure 1:**
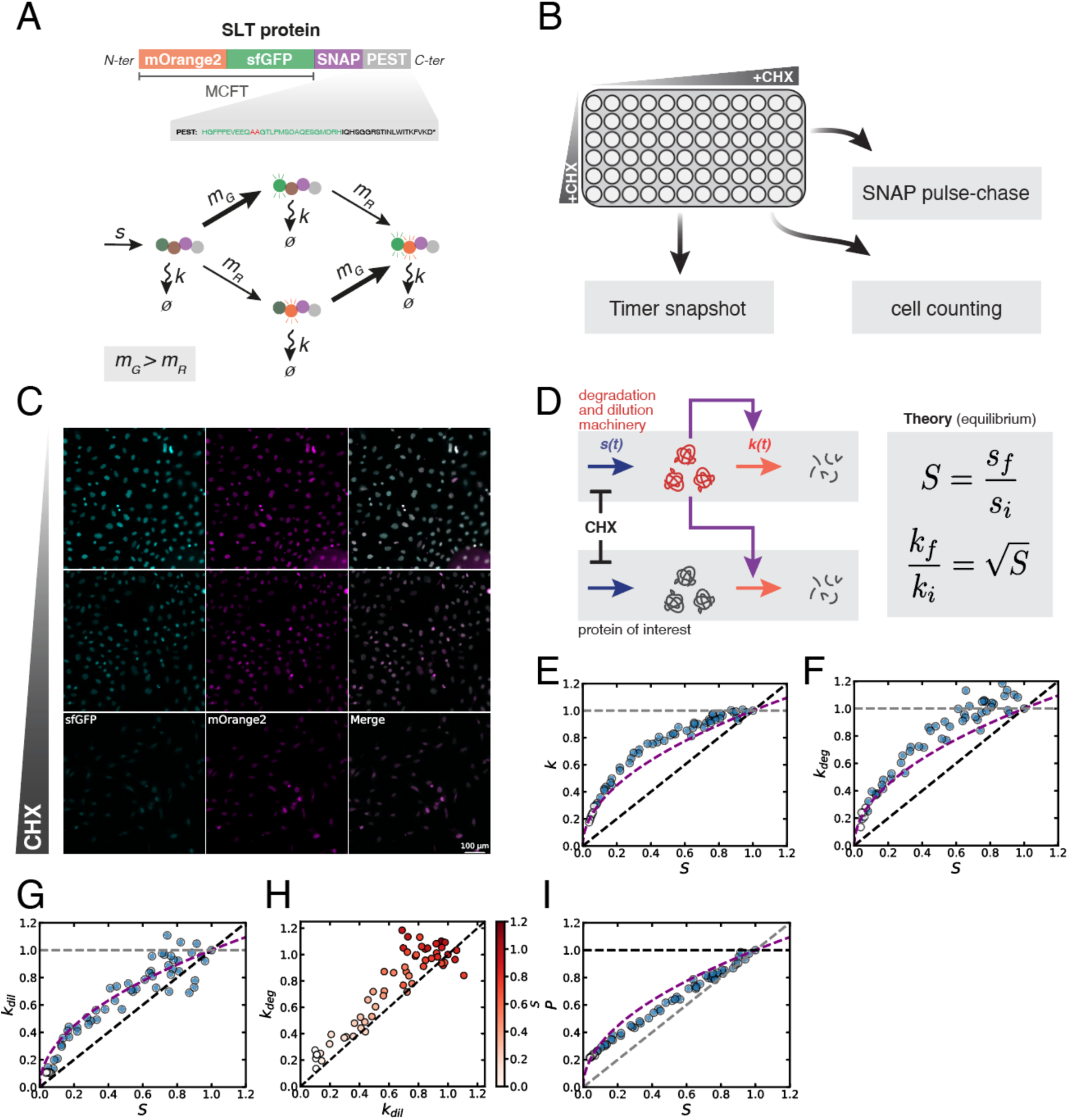
Relationships between *S*, *P*, *k_deg_*, and *k_dil_* upon CHX treatment follow a passive adaptation model. (A) Scheme of the short-lived timer (SLT) construct. MCFT: Mammalian Cell-optimized Fluorescent Timer. (B) Experimental design to determine changes in *S*, *k_deg_, k_dil_* and *P* after treatment with different CHX concentrations. (C) Green (cyan) and Red (magenta) fluorescence snapshots of NIH/3T3 cell populations treated with different CHX concentrations (from top to bottom: 0.008, 0.1125, and 0.5 µg/mL) for 48 hours. (D) Chemical scheme of the passive adaptation model and predicted relationship of *k* and *S* at equilibrium. (E-G) Fold-change in *k* (E), *k_deg_* (F), and *k_dil_* (G) versus fold-change in *S*. (H) Fold-change in *k_deg_* versus fold-change in *k_dil_*. Color bar: fold-change in *S*. x=y diagonal black dashed line: equal fold-change in degradation and dilution rates. (I) Fold-change in *P* versus fold-change in *S*. White dots: data points for which exponential division was lost. Black dashed lines: prediction for perfect adaptation; Gray dashed lines: prediction for no adaptation. Purple dashed curved lines: prediction for passive adaptation. The values shown for *S*, *P*, *k*, *k_deg_*, and *k_dil_* are normalized on the respective values for control conditions.

We then asked how protein decay rates *k* adapt to changes in global protein synthesis rates *S*. Note that *S* itself depends on the availability of the translation machinery, translation efficiency, and the pool of available mRNAs. To allow for global, tunable, fast, and reversible inhibition of global protein synthesis at the level of translation, we decided to use cycloheximide (CHX)^36,37^, which blocks the elongation of polypeptide chains by the ribosome. We treated SLT-expressing NIH/3T3 fibroblasts with 56 different concentrations of CHX ranging from 0.002 to 0.5 µg/ml for 48 hours (Fig.1B, Supplementary Table 1) to fine-tune global translational activity. Crucially, we focused our interpretation on a CHX concentration range at which cells keep proliferating exponentially (up to 0.2 µg/ml, Fig.S1C, STAR Methods) to avoid biases caused by altered cellular health. We then performed SNAP pulse-chase labeling followed by live-cell imaging for 12 hours while maintaining the respective CHX concentrations (Fig.1C). We used sfGFP and mOrange2 fluorescence intensities to infer changes in *S* and *k* (STAR Methods), and the integrated sfGFP intensity to compute the total MCFT protein amount *P*. We also measured *k_deg_* directly by quantifying the SNAP-SiR647 signal decay (Fig.S1D), and protein dilution rates (*k_dil_*) by quantifying changes in cell numbers (Fig.S1E). We used the SNAP-SiR647 pulse-chase experiment to quantitatively calibrate the MCFT (Fig.S1F). As expected, *S* decreased with increasing CHX concentrations (Fig.S1G). *k_deg_*, *k_dil_,* and *P* decreased with increasing CHX concentrations (Fig.S1H-J), indicating that the protein degradation and cell cycle machineries are sensitive to changes in *S*. Results obtained using SNAP pulse-chase labeling and treatment with different concentrations of another protein synthesis inhibitor (anisomycin)^37^ yielded similar results (Fig.S1D, S1K-M, Supplementary Table 2). Altogether, these results indicate that *k* adapts partially to changes in *S*.

We then asked whether decreased production of protein degradation and dilution machineries caused by a decrease in *S* would be sufficient to explain the partial adaptation of *k* to *S* in NIH/3T3 cells. Briefly, we modeled three interdependent sets of proteins: the degradation machinery (DeM), the dilution machinery (DiM), and a protein of interest (POI) (Fig.1D). The degradation and dilution machineries are required for their own decay and the decay of the POI. The model assumes that translation inhibition by CHX scales down *S* of the DeM, DiM, and the POI by the same factor (Fig.1D). As CHX reduces DeM and DiM synthesis, their protein level decreases, subsequently reducing their decay rate. After reaching equilibrium, the protein level of DeM and DiM, and thus the POI decay rate scale as the square root of *S*, leading to the same scaling for the POI (Fig.1D, Supplementary Text). We used our SLT data to ask how well our model explains changes in the activity of the degradation and dilution machineries. The passive adaptation recapitulated the relationships of *S* with *k*, *k_deg_* and *k_dil_* (Fig.1E-G), even though *k_deg_* stayed more slightly stable than expected upon moderate changes in *S* (Fig.1F and 1H). This suggests that spare degradation resources allow to maintain *k_deg_* upon a modest decrease in *S*. In addition, *k_dil_* adapted more than expected at very low *S* (<0.3), which might be explained by altered cell viability (as assessed by loss of exponential proliferation) in this regime. We also found that changes in *P* upon changes in *S* could be approximated by the passive adaptation model (Fig.1I). Log-likelihood computation and bootstrapping for model selection (STAR Methods) confirmed these observations (Fig.S1N-O).

We next asked whether changes in *k_deg_* and *k_dil_* upon a decrease in *S* are interdependent. Since the proteasome is a long-lived machinery^38–41^, we reasoned that in dividing cells, its concentration might be more dependent on *k_dil_* than *k_deg_*. To test this hypothesis, we took advantage of the global gene expression and cell proliferation regulation by the MYC transcription factor^42^ and the availability of a specific MYC inhibitor 10058-F4^43^ (MYCi). Since MYC enhances the global transcriptional and translational activity and directly activates cell cycle regulators to stimulate cell proliferation^42^, we reasoned that MYCi would both decrease *S* and *k_dil_* independently. We treated NIH/3T3 cells for 48 hours with 56 doses of MYCi ranging from 0.26 to 64 µM (Fig.S1P, Supplementary Table 3). As expected, we observed a MYCi dose-dependent decrease in *S* (Fig.S1P-R) and a strong, close-to-perfect adaptation of *k_dil_* to changes in *S* (Fig.S1S-T). In contrast, *k_deg_* stayed constant until *S* went below 50% of its value in control conditions (Fig.S1U-W). This suggests that a strong decrease in *k_dil_* may abolish changes in *k_deg_* in response to a decrease in *S*, and thereby, that *k_deg_* is dependent on *k_dil_*.

### A passive adaptation model explains changes in *P* and *k* for other proteins and cell types

To test the generality of the passive adaptation model, we generated NIH/3T3 cell lines constitutively expressing the MCFT-SNAP with a mutated, non-functional PEST (long-lived timer or LLT), or fused to the NANOG, ESRRB, or SOX2 proteins. This resulted in MCFT-labeled proteins characterized by different protein-specific *k_deg_* (ps*k_deg_*) at steady state (Fig.S2A) and thus distinct contributions of *k_deg_* and *k_dil_* to *k* (Fig.S2B). Given that the LLT has a very low ps*k_deg_* (Fig.S2A), CHX treatment was extended to 6 days to ensure reaching a steady state. The passive adaptation model recapitulated well the changes in *k* upon CHX treatment for all tested proteins (Fig.2A-D). To quantify changes of total protein levels, we treated cells with different CHX concentrations for 6 days, labeled all intracellular proteins using N-hydroxysuccinimidyl (NHS) ester that reacts with free amino groups^30^, and quantified changes in the nuclear NHS ester signal (Fig.S2C and Fig.2E). The passive adaptation model explained well the changes in total nuclear protein content at moderate changes in *S*. When reaching very low *S* levels, changes in total protein levels started to deviate from the passive adaptation model. This is expected given the long median half-life of proteins in NIH/3T3 cells (16.7 ± 6.2 (median ± std) hours from ^44^, Supplementary text) that implies a dominant impact of *k_dil_* over *k_deg_*.

**Figure 2:**
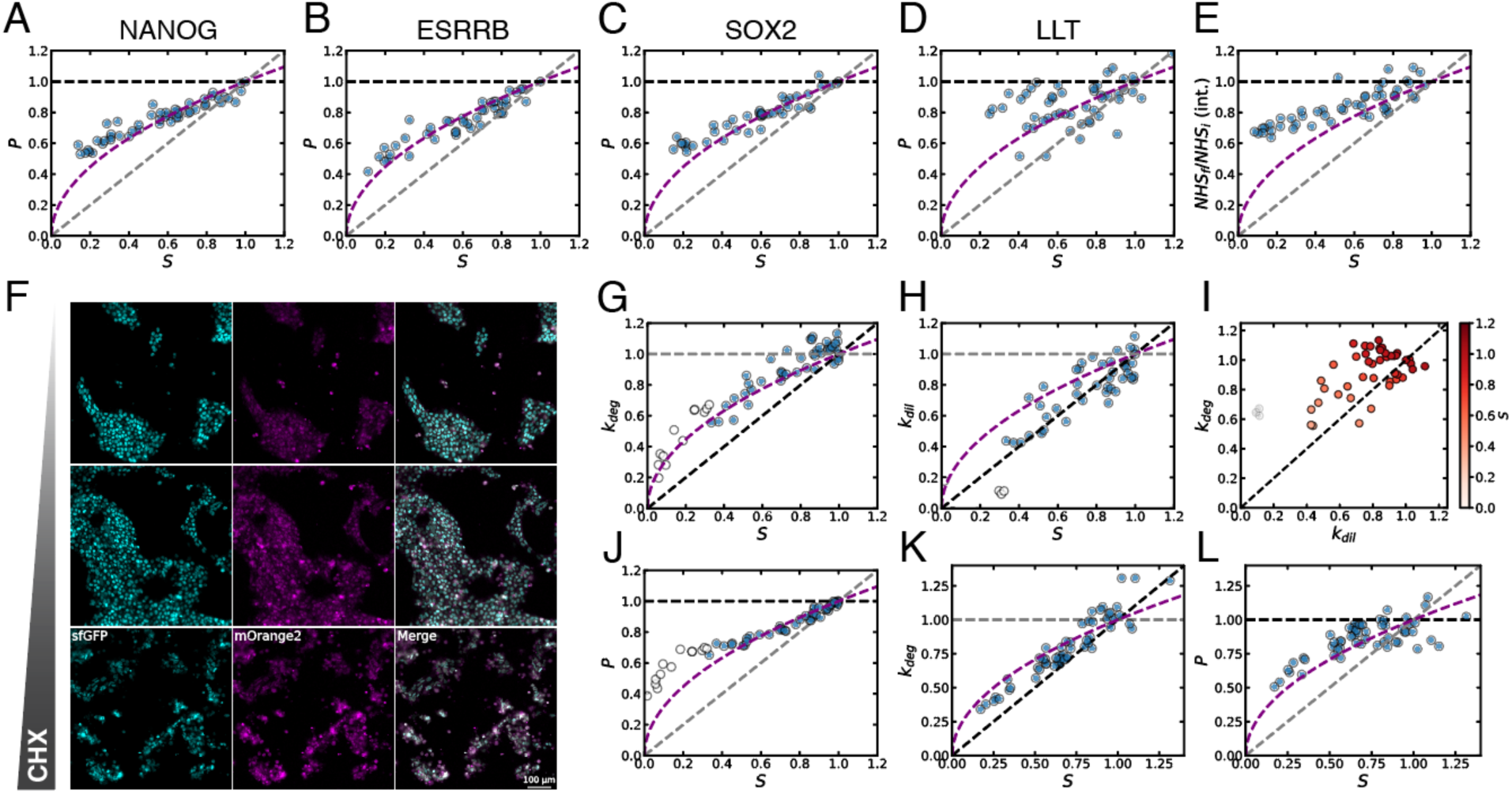
Passive adaptation of *k* and *P* to *S* for other proteins, and for SLT in hESCs and hESC-derived astrocytes. (A-D) Fold-change of *P* with respect to *S* for NANOG (A), ESRRB (B), SOX2 (C), and the LLT (D). (E) Fold-change in total nuclear protein levels with respect to *S*, measured with NHS-ester. (F) Green (cyan) and red (magenta) fluorescence snapshots of hESC SLT cell populations treated with different CHX concentrations (from top to bottom: 0.008, 0.1125, and 0.5 µg/mL) for 48 hours. (G-H) Fold-change in *k_deg_* (G) and *k_dil_* (H) vs fold-change in *S* in hESCs. (I) Fold-change in *k_deg_* with respect to the change in *k_dil_* in hESCs treated with CHX. Color bar: fold-change in *S* in each condition. (J) Fold-change in *P* versus *S* for CHX treatment in hESCs. (K) Fold-change in *k_deg_* vs fold-change in *S* for CHX treatment in human astrocyte-enriched culture. (L) Fold-change in *P* versus *S* for CHX treatment in human astrocyte-enriched culture. (A-E, G-H, J-L) Black dashed lines: prediction for perfect adaptation; Gray dashed lines: prediction for no adaptation. Purple dashed curved lines: prediction for passive adaptation. White dots: data points for which exponential division was lost. The values shown for *S*, *P*, *k_deg_* and *k_dil_* are normalized on the respective values for control conditions.

We then asked if passive adaptation could also explain the adaptation of *k* to *S* in other mammalian cell types. We generated a knock-in human embryonic stem cell line (hESC) in which the SLT was integrated into the CLYBL locus^45^ (STAR Methods, Fig.S2D-F). Upon treatment with different doses of CHX (Fig.2F), *k_deg_* followed the passive adaptation model (Fig.2G), while *k_dil_* adapted slightly more efficiently (Fig.2H-I). Changes in *P* were well-approximated by the passive adaptation model (Fig.2J).

We then asked how protein decay adapts to changes in protein synthesis rates in differentiated, non-dividing cells. To do so, we differentiated SLT-expressing hESCs into astrocytes^46^ (STAR Methods) (Fig.S2G-J). Importantly, most cells were negative for the proliferation marker Ki67, indicating their post-mitotic state (Fig.S2G-H). We then treated SLT-expressing astrocyte-enriched cultures with different concentrations of CHX (Fig.S2K-L), and found that *k*, and *P* followed the passive adaption model (Fig.2K-L). Together, these data suggest that passive adaptation of the decay rate *k* to the synthesis rate *S* also occurs in the absence of cell division in differentiated post-mitotic cells.

### *k* adapts to *S* within 10 hours

We then developed a superstatistical Bayesian approach^47–49^ to infer time variations of *S* and *k* from MCFT traces (Methods, Supplementary Text, Fig.S3A). To validate our inference method, we generated synthetic sfGFP and mOrange2 fluorescence trajectories (Fig.S3B-C) imposing defined variations of *S* and *k*. Our inference robustly recovered ground truth time variations of *S* and *k* from the fluorescence trajectories (Fig.S3D-E).

We next used a cell line in which MCFT expression is controlled by a doxycycline (dox)-inducible promoter in NIH/3T3 cells^31^, allowing the alteration of the synthesis rate of the MCFT only, without expected changes in its ps*k_deg_*. We used 28 different step changes in dox concentrations (Fig.3A) and quantified sfGFP and mOrange2 fluorescence over time (Fig.S3F). In all cases, we correctly inferred expected changes in *S* and a mostly unaltered *k* (Fig.3A). Furthermore, data retrodiction (Methods, Supplementary Text) confirmed that our inference accurately reproduces the input data (Fig.S3G). We conclude that our inference scheme allows robust discrimination between changes in *S* and *k* from time traces of the MCFT.

**Figure 3:**
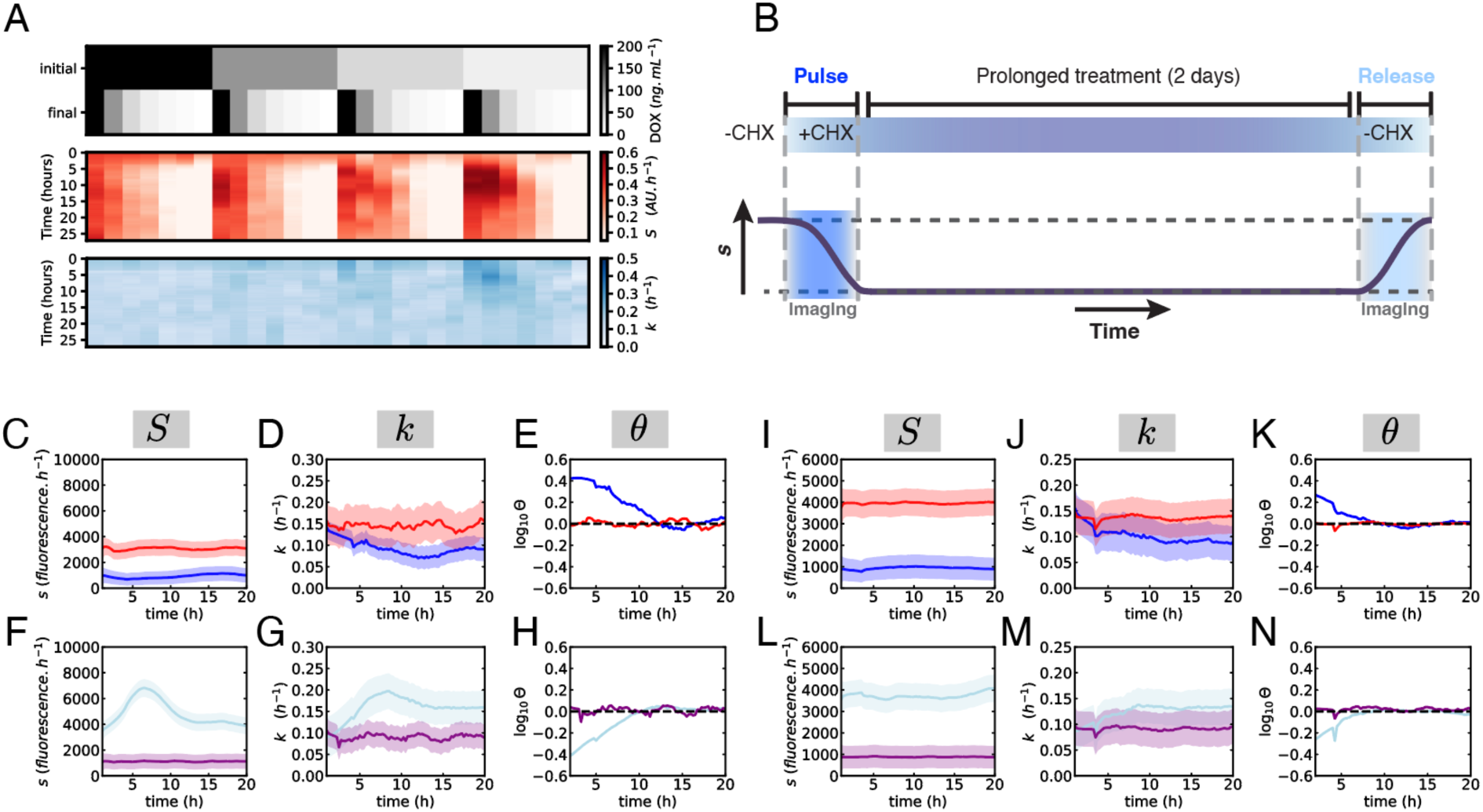
Passive adaptation occurs with a time scale of 5-15 hours. (A) Upper panel: dox concentration changes between initial and final concentrations used just before imaging. Heatmaps: temporal variations of *S* (red) and *k* (blue) inferred from SLT trajectories. (B) Experimental design of CHX pulse and release experiments. (C-N) Time-lapse imaging for NIH/3T3 (C-H) and hESCs (I-N). *S* (C,I) and *k* (D,J) trajectories inferred from MCFT measurements during CHX pulse. Red: control conditions. Dark blue: CHX pulse. Purple: continuous CHX treatment. Light blue: CHX release. (E,K) Time variation of the imbalance for the CHX pulse. *S* (F,L) and *k* (G,M) trajectories inferred from MCFT measurements during CHX release. (H,N) Time variations of the imbalance for CHX release. (C-G) and (I-M), plain lines: average of the posterior distribution; shaded regions: SD of the posterior distribution.

We then aimed at quantifying the time scales characterizing the adaptation of *k* to alterations in *S*. To do so, we performed time-lapse imaging of the MCFT in NIH/3T3 constitutively expressing the SLT before, during treatment with 0.1 µg/mL of CHX and after CHX release (Fig.3B, Supplementary Movies 1-2). We then used our inference algorithm to determine changes in *S* and *k* from SLT fluorescence time traces. Upon CHX treatment, *S* sharply decreased (Fig.3C) while *k* progressively decreased until reaching a lower plateau (Fig.3D). To quantify the adaptation time delay of *k* to *S*, we used a metric (*θ*) that we recently developed, which allows quantifying the imbalance between protein influx (governed by *S*) and outflux (governed by *k*)^50^. Briefly, the imbalance reads *θ = Pk/S*, such that at equilibrium *θ = 1*. *θ > 1* or *< 1* indicates an excess of protein outflux or influx, respectively^50^. *θ* reached its steady-state value around 10 hours after CHX addition (Fig.3E). Upon CHX release, we observed similar dynamics and delay in the adaptation of *k* to *S* and in *θ* (Fig.3F-H). We performed the same CHX pulse and release experiments in hESCs (Supplementary Movies 3-4), which resulted in a similar delay in the adaptation of *k* to *S* (Fig.3I-N). Finally, we also performed MYCi pulse and release experiments in NIH-3T3, and we found that *S* and *k* reached a steady state within a similar time frame (Fig.S3H-M).

### Naive mouse embryonic stem cells display a distinct, biphasic adaptation mode

Next, we aimed to characterize the adaptation of *k* and *P* to *S* in naive mouse embryonic stem cells (mESCs). mESCs display a modified cell cycle with an altered G1-S checkpoint and an inactive Rb protein^51^. Rb allows coordination of cell growth – itself dependent on protein content accumulation – and cell division^52^. We thus reasoned that in mESCs, *k_dil_* may react differently to changes in *S* as compared to NIH/3T3 and hESCs. We generated an mESC line in which the SLT is knocked into the ROSA26 locus^53^ (STAR Methods, Fig.S4A-C) and quantified the response of *k* and *P* to changes in *S* (Fig.4A-E). In contrast to NIH/3T3 and hESCs, *k_deg_* over-adapted to changes in *S* (Fig.4B), and *k_dil_* remained high until *S* dropped below *circa* 60% of its maximal value (Fig.4C-D). Both the SLT and NHS-ester-stained nuclear proteome remained virtually unchanged upon a decrease in *S*, underscoring the close-to-perfect adaptation of *k* to *S* (Fig.4E-F, Fig.S4D). We then performed time-lapse imaging to monitor changes in *k* upon a 0.1 µg/ml CHX pulse (Supplementary Movie 5) and release (Supplementary Movie 6). *S* initially dropped upon CHX addition but rapidly increased thereafter before dropping again to reach a slightly higher value than immediately after CHX addition (Fig.4G). *k* was decreased immediately at the beginning of imaging and stabilized after about 10 hours (Fig.4H-I). This suggests that an active, fast-acting mechanism allows to partially counteract CHX inhibition and to rapidly shut down protein degradation upon decrease of *S*. Upon CHX release, we observed a rapid increase in *S* that then dropped again, before increasing to levels of control conditions (Fig.4J), while *k* showed a rapid initial increase and reached steady state towards the end of the imaging period (∼20h) (Fig.4K-L). Therefore, mESCs adopt a biphasic adaptation mode in which *k_deg_* and *S* respond to CHX within a few hours, but full adaptation occurs in a slower second phase.

**Figure 4:**
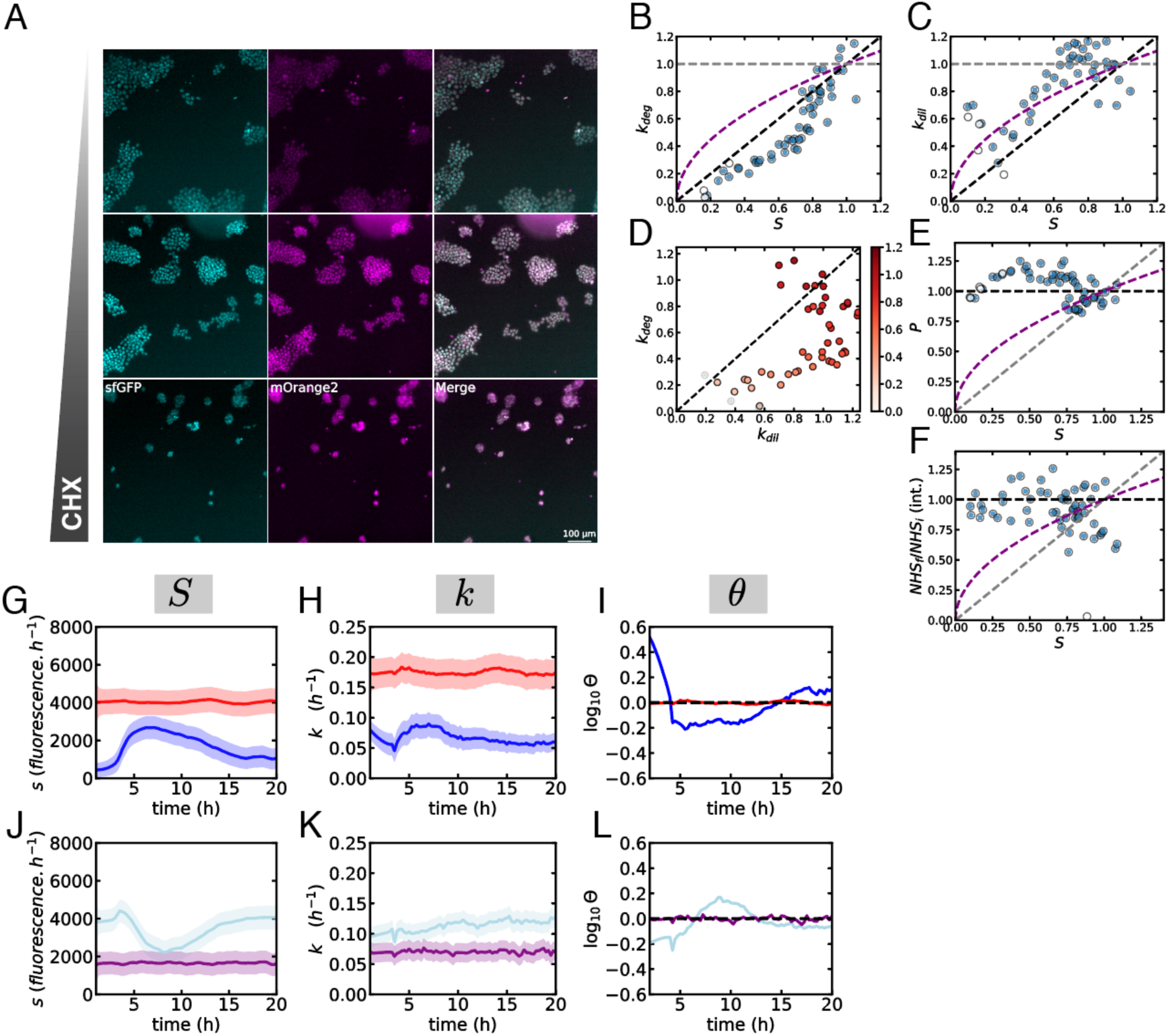
mESC adaptation is inconsistent with a purely passive mechanism. (A) Green (cyan) and red (magenta) fluorescence snapshots of mESC SLT cell populations treated with different CHX concentrations (from top to bottom: 0.008, 0.1125, and 0.5 µg/mL) for 48 hours. (B-C) Changes in *k_deg_* (B) and *k_dil_* (C) vs *S* for CHX treatment in mESCs. (D) Changes in *k_deg_* vs *k_dil_* in mESCs treated with CHX. Color bar: fold-change in *S*. (E) Changes in *P* vs *S* for the SLT. (F) Fold-change in total nuclear *P* versus *S*, measured with NHS-ester. (B,C,E,F) Black dashed lines: prediction for perfect adaptation; Gray dashed lines: prediction for no adaptation. Purple dashed curved lines: prediction for passive adaptation. (B-F) The values shown for *S*, *P*, *k_deg_* and *k_dil_* are normalized on the respective values for control conditions. (G,H) *S* (G) and *k* (H) trajectories inferred from MCFT measurements during CHX pulse. (I) Time variation of the imbalance for CHX pulse. (J-K), *S* (J) and *k* (K) trajectories inferred from MCFT measurement during CHX release. (L) Time variation of the imbalance for CHX release. Red: control conditions. Dark blue: CHX pulse. Purple: continuous CHX treatment. Light blue: CHX release. For G,H,J,K, the average (plain line) and the SD (shaded region) of the posterior distribution are represented for each timepoint.

### Physiological variability in protein turnover recapitulates cell type-specific adaptation

To ensure that the adaptation we uncovered is not caused by unspecific effects of translation inhibitors and reflects physiological changes of *k* in response to fluctuations in *S*, we took advantage of the natural fluctuations of *S* in unperturbed conditions^31^ and determined the relationship between *S*, *k,* and *P* at the single cell level. We used time-lapse imaging data acquired in the absence of perturbation in i) 5 NIH/3T3 cell lines expressing the SLT, LLT, and the MCFT fused to NANOG, ESRRB, and SOX2, respectively; ii) 40 mESC lines in which the MCFT is internally fused to 40 different endogenously expressed proteins^31^. We have previously shown that the majority of these 40 proteins exhibit only minor fluctuations in *S*, *k*, and protein concentration [*P*] over the cell cycle in single cells^31^. This quasi-steady state allows us to compute *k* for single cells as before for population averages (STAR Methods). Similarly, we computed *S_conc._* and [*P*] for single cells. Note that *S_conc._* is a per-cell protein synthesis rate in concentration per hour unit (STAR Methods). Since *S_conc._* does not correlate with nucleus size (Fig.S5), it can be considered a proxy of single-cell protein synthesis rate, *S* being proportional to its population average. We then normalized the data from all proteins to their median value for *S_conc_*, *k,* and [*P*], and aggregated all data points. In NIH/3T3 cells, the correlations of *k* and [*P*] with *S_conc_* followed the passive adaptation model (Fig.5A-B). In contrast, in mESCs the correlation between *S_conc_* vs *k* and [*P*] was closer to a perfect adaptation (Fig.5C-D). Altogether, our data demonstrate that cell type-specific adaptation modes of *k* to *S* are operating upon physiological fluctuations of *S*.

**Figure 5:**
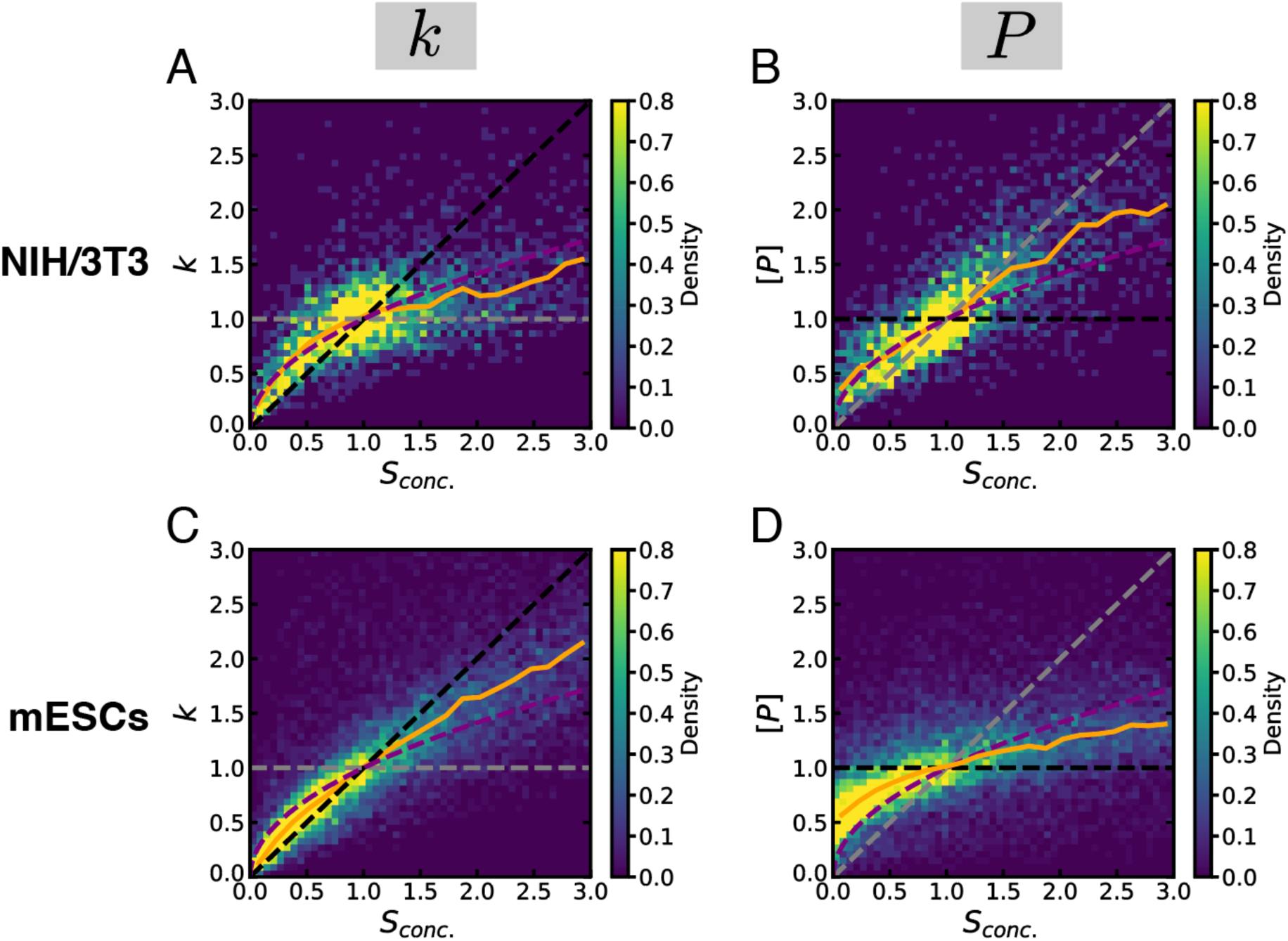
Adaptation of *k* and [*P*] *to S_conc_* in unperturbed conditions. (A-B) *k* (A) and [*P*] (B) versus *S_conc_* in NIH/3T3 cells. (C-D) *k* (C) and [*P*] versus *S_conc_* (D) in mESCs. 2D histograms for aggregated and normalized data for 5 (NIH/3T3) or 40 (mESCs) cell lines are shown. N > 4,000 (NIH/3T3) or >10,000 (mESC) cells. Black dashed lines: prediction for perfect adaptation; Gray dashed lines: prediction for no adaptation. Purple dashed curved lines: prediction for passive adaptation. Orange plain line: binned median of the aggregated data.

### Sustained changes in mTOR signaling drive robust adaptation in mESCs

Changes in global translation rates can modulate mTOR activity by altering the intracellular concentration of free amino acids^36,37,54–58^. Since mTOR activity regulates both *S* and *k*, we asked whether changes in mTOR activity contribute to the adaptation of *k* to *S*. We first quantified mTOR activity by measuring phosphorylation of the mTOR target p70S6K after 1h, 2h, 5h, and 48h of 0.1μg/mL CHX treatment by ELISA assay in NIH/3T3 cells and mESCs. mTOR activity acutely increased and peaked after 2h in both cell types, before slowly decreasing, to a lesser extent in mESCs as compared to NIH/3T3 cells (Fig.6A). We next asked whether changes in ribosomal protein gene expression in mESCs, which are targets of mTORC1^59^, could explain the rebound in *S* and its elevated levels after circa 20h of CHX compared to the earliest time points (Fig.4G). We performed RNA-seq at different timepoints (0, 1, 2, 5, 48 hours) after CHX pulse. Within the first hours of CHX treatment, ribosomal protein genes were markedly upregulated in mESCs but more modestly in NIH/3T3 cells, and after 48 hours they decreased below their initial levels in NIH/3T3 but not mESCs (Fig.S6A-C). Other genes involved in protein turnover (proteasomal and mitochondrial genes) followed the same dynamics but with a smaller amplitude. Taken together, this suggests that mESCs exhibit a stronger and more sustained adaptation of mTOR signaling and expression of ribosomal protein genes than NIH/3T3 cells.

**Figure 6:**
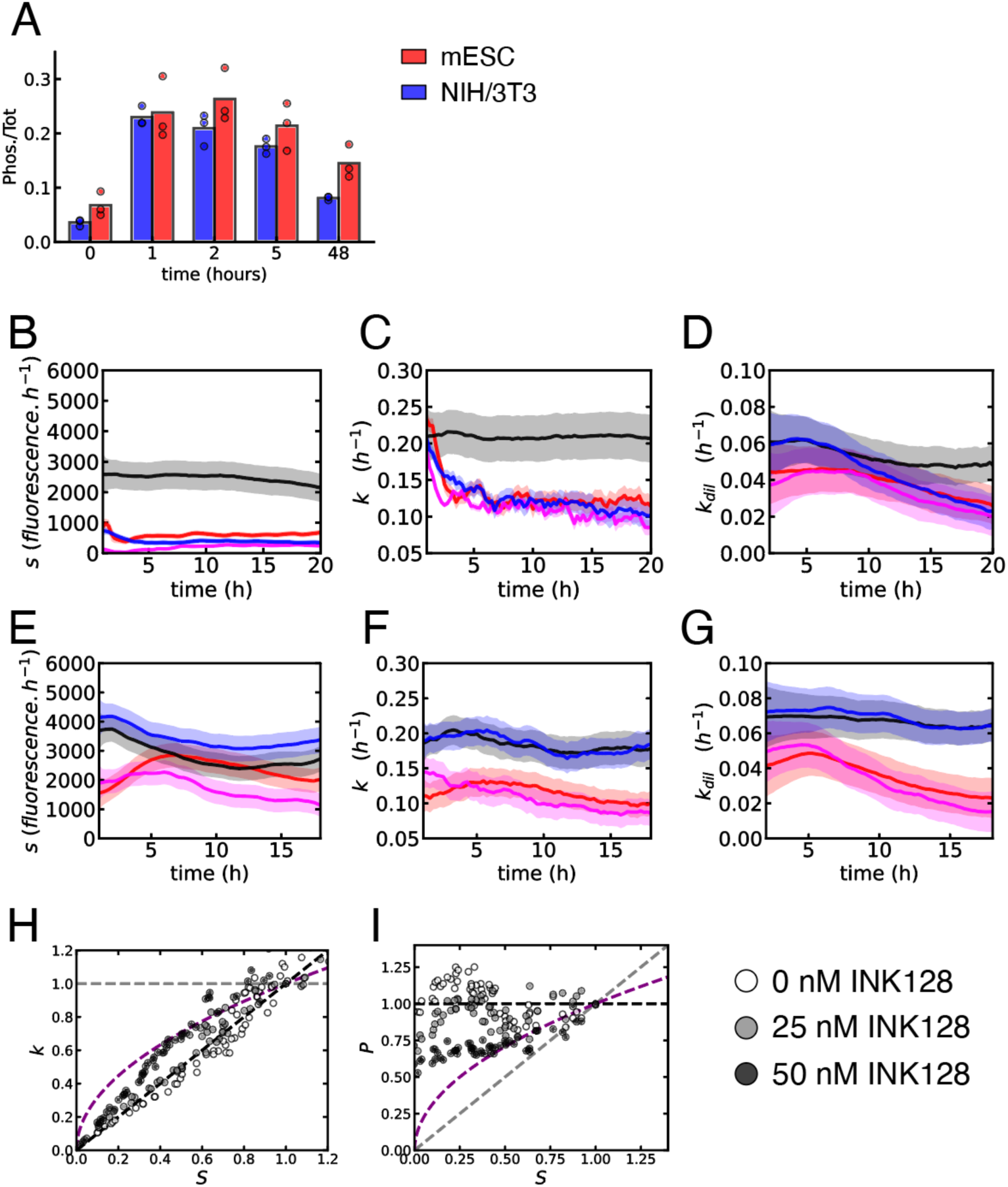
mTOR-dependence of the mESC response to protein synthesis inhibition. (A) Phosphorylation kinetics of p70S6K measured by ELISA after CHX treatment (0.1 µg/mL) in mESC and NIH/3T3. (B-D) *S* (B), *k* (C), and *k_dil_* (D) trajectories inferred from MCFT (*S* and *k*) and cell number (*k_dil_*) measurements during CHX pulse for NIH/3T3 cells. (E-G) *S* (E), *k* (F), and *k_dil_* (G) trajectories inferred from MCFT (*S* and *k*) and cell number (*k_dil_*) measurements during CHX pulse for mES cells. Black: control conditions. Red: CHX treatment. Blue: 25nM INK128 treatment. Pink: CHX + 25 nM INK128 treatment. The average (plain line) and the SD (shaded region) of the posterior distribution are represented for each timepoint. (H-I) Fold-change in *k* (H) or *P* (I) versus fold-change in *S* for mES cells long-term treatment with different CHX concentration in the presence of 0, 25, or 50 nM of INK128. Black dashed lines: prediction for perfect adaptation. Gray dashed lines: prediction for no adaptation. Purple dashed curved lines: prediction for passive adaptation.

We next performed CHX pulse time-lapse imaging in mESCs and NIH/3T3 with and without the mTOR inhibitor INK128 or the ISR inhibitor ISRIB. Cells were pre-treated with either inhibitor for 1 h prior to imaging. In both mESCs and NIH/3T3 cells, 200 nM ISRIB led to slightly increased *S*, but the response of *k* was similar to the one of untreated cells, indicating that the ISR is not required for the adaptation of *k* to *S* (Fig.S6D-G). Since a full inhibition of the mTOR pathway causes mESCs to stop proliferating and enter diapause^60,61^, we used a low INK128 dose (25nM) that allowed mESCs to maintain their proliferation rate (Fig.S6H). In NIH/3T3 cells, INK128 alone markedly decreased *S,* and *k* adapted to changes in *S* following the passive adaptation model (Fig.6B-D and Fig.S6I). When combined with CHX, INK128 had almost no impact on the dynamic changes in *S* and *k* for up to 18 hours (Fig.6B-D). This suggests that in NIH/3T3 cells, mTOR activity is important to sustain *S* but is not involved in the adaptation of *k* to changes in *S*. In mESCs, INK128 had virtually no impact on *S* in the absence of CHX (Fig.6E-G). However, the rapid initial boost in *S* and decrease in *k_deg_* caused by CHX was impaired by mTOR inhibition. In the longer term (>15 hours), mTOR inhibition in the presence of CHX resulted in strongly decreased *S* levels compared with CHX alone and underadaptation of *k* to changes in *S* (Fig.6E-G, Fig.S6J). Since INK128 did not impact *k_dil_* in mESCs (Fig.S6H), this suggests that mTOR activity directly contributes to the adaptation of *k_deg_* to *S* in mESCs.

We then repeated the CHX plate experiment in mESCs in the presence of either 25 or 50nM of INK128. Remarkably, INK128 weakened the adaptation of *k* to *S* in a dose-dependent manner, closely mimicking passive adaptation at 50nM until *S* = 0.3, matching regimes below which exponential proliferation becomes less robust (Fig.6H-I and Fig.4B-C). Altogether, our results indicate that mESCs complement passive adaptation by a sustained adjustment of mTOR-signaling, resulting in robust adaptation of *k_deg_* to *S*.

### Proteome composition and dynamics upon passive and mTOR-driven adaptation

We next quantified long-term changes in the proteome of mESCs and NIH/3T3 upon CHX treatment. We cultured NIH/3T3 cells and mESCs for 4 and 2 days, respectively, with or without 0.1 µg/ml of CHX. We chose these respective CHX treatment durations to maximize the likelihood of reaching a new steady state for all proteins, based on the respective cell cycle durations of NIH/3T3 (ca 22h) and mESCs (ca 12h). We then extracted proteins and performed label-free quantification (LFQ) mass spectrometry (STAR Methods). While 10% and 2.8% of proteins were down- or up-regulated in NIH/3T3 (Fig.7A-B), respectively, only 0.6% and 0.1% were down- or up-regulated in mESCs (Fig.7C-D). Importantly, at this CHX concentration the immediate impact on *S* levels was even stronger in mESCs than in NIH/3T3 cells (earliest time points in Fig.3C and Fig.4G), suggesting that mESCs have a superior ability to maintain their protein levels upon perturbation of *S*. Ribosomal proteins were well-preserved in both cell types, but proteins involved in mitochondrial or proteasomal functions were slightly downregulated in NIH/3T3 cells but not in mESCs (Fig.S7A-C), in line with passive adaptation of cellular resources governing protein decay in NIH/3T3.

**Figure 7:**
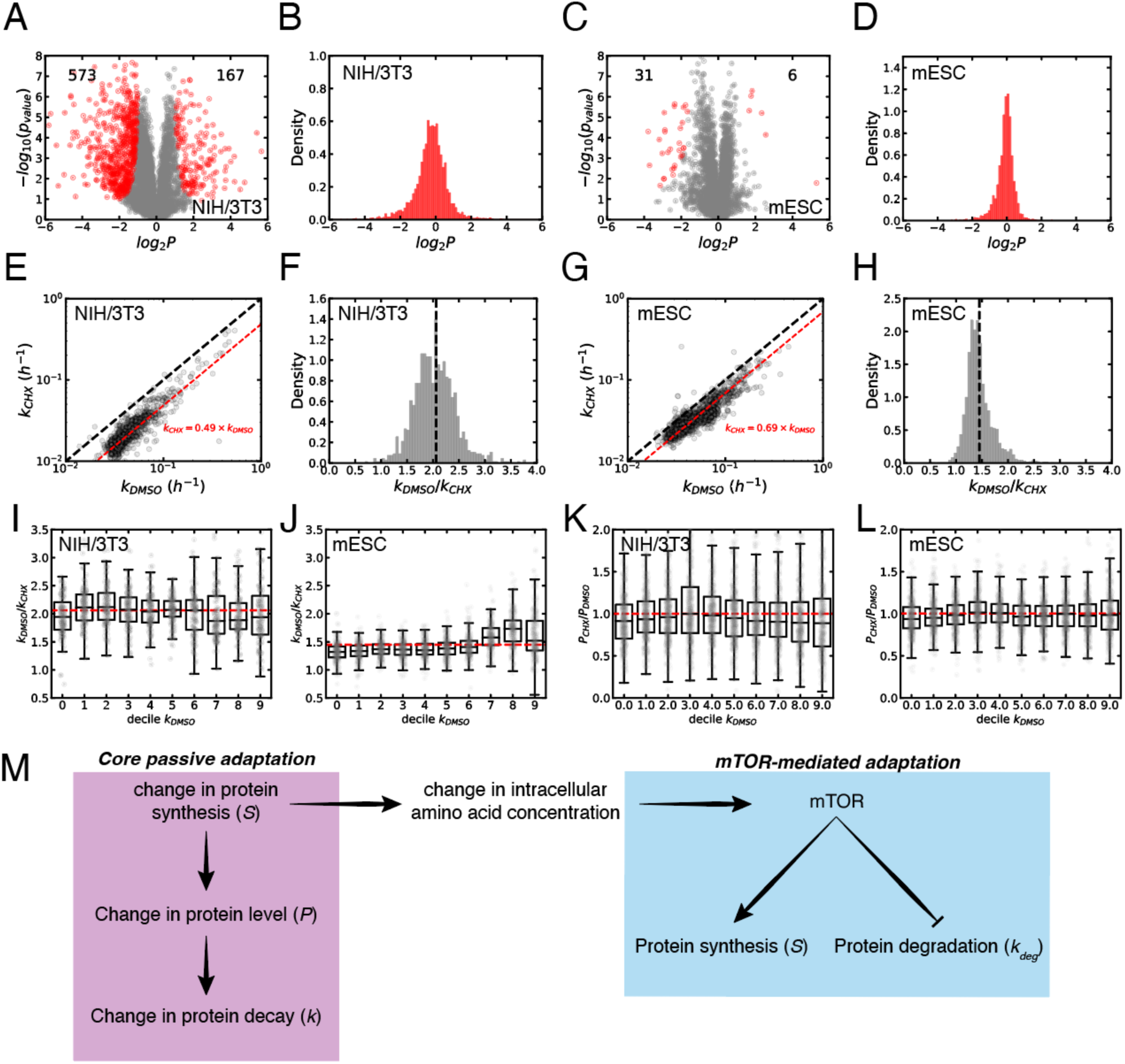
Proteome-wide impact of a prolonged decrease in *S*. (A-D) Label-free proteome quantification for NIH/3T3 cells (A-B) or mESCs (C-D) treated with CHX, leading to the identification of 5875 and 4503 proteins, respectively. (A, C) The average of four replicates is plotted for all identified proteins. Red dots: proteins with significant (FDR = 0.01) fold-change in their levels between CHX and control conditions. Each point is the average over quadruplicates of the protein-specific level fold-change. (B, D) Distribution of protein level fold-changes between CHX and control conditions. (E,G) Decay rate in (0.05 µg/mL) CHX versus in DMSO condition for proteins measured in dynamic SILAC (STAR Methods and Fig.S8), for NIH/3T3 (E) and mESCs (G). The black dotted line represents *k_CHX_ = k_DMSO_*. The red dotted line represents the mean fold-change in decay rate between CHX and DMSO conditions. (F,H) Density of the ratio of the decay rates computed in DMSO, *k_DMSO_*, and CHX, *k_CHX_*, conditions for unique proteins in NIH/3T3 (F) and mESCs (H). (I-J) Measured decay rate fold-change between CHX and control conditions, for NIH/3T3 (I) and mESC (J), binned according to ps*k_deg_* in DMSO condition. (K-L) Measured protein level fold-change between CHX and control conditions, for NIH/3T3 (K) and mESC (L), binned according to ps*k_deg_* in DMSO condition. (M) Model of core passive and facultative mTOR-mediated coordination of protein synthesis and degradation.

To determine the proteome-wide adaptation of *k* to changes in *S*, we performed dynamic Stable Isotope Labeling by Amino acids in Cell culture (dSILAC^19,44,62–64^) (STAR Methods, Supplementary Text, and Fig. S7D) in the presence or absence of 0.05 μg/mL of CHX during 72 hours, for both NIH/3T3 and mESCs. We used a CHX concentration of 0.05 μg/mL to avoid total arrest of cell growth for NIH/3T3 cells cultured in the SILAC DMEM medium, which lacks growth factors (STAR Methods). Cellular proteins were initially labeled using a medium containing light (NIH/3T3) or medium (mESC) arginine and lysine amino acids, which was then exchanged for a medium containing heavy (NIH/3T3) arginine and lysine. Samples were collected 0h, 1h, 2h, 4h, 6h, 12h, and 24h (plus 48h for NIH/3T3) after medium exchange. After protein extraction, sample preparation, and mass spectrometry, data were fitted to compute the decay rate per protein (STAR Methods). The decay rate was determined for 2050 (DMSO) and 2629 (CHX) proteins in NIH/3T3, and 3102 (DMSO) and 3380 (CHX) proteins in mESCs. The *k* we inferred for NIH/3T3 proteins correlated strongly with previous NIH/3T3 dSILAC data^44^ (Fig.S7E). Upon CHX treatment, virtually all proteins exhibited a decreased *k* in both NIH/3T3 and mESCs (Fig.7E-H and Fig. S7F-G), confirming the proteome-wide impact of changes in *S* on *k*. Importantly, LFQ analysis of protein levels using the 0h time point confirmed that also for identical CHX treatment duration, NIH/3T3 displayed larger changes in protein levels than mESCs (Fig.S7H-I).

We next reasoned that a quantitatively different adaptation of *k_dil_* vs *k_deg_* in NIH/3T3 and mESCs should result in distinct changes of *k* and *P* for proteins with different ps*k_deg_*. In NIH/3T3, *k_dil_* adapted slightly better than *k_deg_*, while the opposite was true in mESCs. This implies that in NIH/3T3, *k_deg_*-dominated proteins (high ps*k_deg_*) should experience a weaker adaptation of *k*. Conversely in mESCs, the *k_dil_*-dominated proteins (low ps*k_deg_*) should experience a weaker adaptation of *k*. Remarkably, changes in *k* upon CHX treatment as a function of the ps*k_deg_* calculated from our dSILAC data in DMSO conditions closely matched these predictions (Fig.7I-J and Fig.S7J-K). In line with these findings, protein levels were preferentially decreased for proteins with high ps*k_deg_* in NIH/3T3 cells but not in mESCs (Fig.7K-L). Therefore, cell type-specific adaptation modes have distinct, predictable consequences on the regulation of proteome homeostasis upon fluctuating protein synthesis rates.

## Discussion

Here we present a novel microscopy and computational approach allowing to uniquely quantify protein synthesis and decay rates from both snapshots and time-lapse imaging data. This enabled us to quantify protein turnover dynamics at an unprecedented temporal resolution, unravelling how protein decay is coordinated with protein synthesis. We demonstrate that mammalian cells systematically adjust protein decay rates to variations in protein synthesis rates. We show that a passive adaptation mechanism allows partial adaptation of protein decay to protein synthesis rates, resulting in imperfect proteome maintenance. An additional mTOR-driven adaptation layer allows naïve mouse pluripotent stem cells to further optimize the adaptation of protein degradation rates to protein synthesis rates, leading to near-perfect maintenance of their proteome (Fig.7M).

Our passive adaptation model hinges on structural relationships between protein synthesis and the protein decay machinery. While changes in active protein degradation strongly depend on the protein dilution rate in dividing cells, non-dividing cells such as astrocytes also modify their protein degradation rate in response to fluctuations in protein synthesis rates in a quantitatively similar way. This suggests that the turnover of their degradation machinery is actively regulated, consistent with observations that non-dividing cells exhibit a higher degradation rate for long-lived proteins than dividing cells^65^. Passive adaptation allows only for partial maintenance of overall protein levels, but the roughly comparable adaptation of *k_deg_* and *k_dil_* in dividing cells mitigates the imbalance between proteins decaying mainly through dilution vs degradation.

In contrast with other cell types, mESCs robustly decrease *k_deg_* and partially rescue translational efficiency upon decrease of translational resources through their ability to sustainably increase mTOR signaling. This additional adaptation layer further improves the resilience of the mESC proteome, resulting in near-perfect maintenance of protein levels when their translational capacity is impaired. Interestingly, mESCs and the inner cell mass of the pre-implantation blastocyst use a unique strategy for their amino acids supply based on extracellular protein uptake^66^. This mechanism may explain the robust viability of the preimplantation embryo even in the absence of exogenous amino acids^67^, which may be complemented by the adaptation mechanisms we describe here to maximize the proteome resilience of the early embryo.

Our work has further methodological and biological implications beyond the model systems we used. First, the rapid and universal adaptation of protein decay to changes in protein synthesis rates directly questions the relevance of the CHX pulse approach to determine protein half-lives^36,68^. Second, two views conflict on the predominant roles of species-specific differences in protein synthesis^69^ versus degradation rates^70,71^ in regulating the pace of somitogenesis. We argue that the fundamental interconnection between protein synthesis and protein degradation rates makes it extremely challenging to causally address the functional relevance of these rates independently. Third, mTOR inhibition was both reported to decrease^29^ and increase^27^ protein degradation rates. We found that increased mTOR activity upon a decrease in translational resources contributes to a further decrease in protein degradation rate, in agreement with^27^. However, the modulation of protein synthesis by mTOR signaling indirectly influences protein degradation rates through passive adaptation, in line with decreased protein degradation rates observed during long-term (>16 hours) mTOR inhibition^29^.

In summary, our study unravels the pivotal resilience mechanisms that mammalian cells leverage to buffer variations of proteome concentrations when confronted with fluctuations in resources governing protein synthesis rates.

## Acknowledgments

This work was supported by EPFL core funding and the Synapsis Foundation Switzerland. The authors want to thank Almut Eisele, Felix Naef, and Maike Hansen for their careful reading of the manuscript and their comments. We would like to thank Louis-Alexandre Ongaro for testing the feasibility of the astrocyte differentiation protocol and performing the initial differentiation to astrocyte progenitors. Fluorescence-activated cell sorting was performed at the EPFL Flow Cytometry Core Facility (EPFL-FCCF). Microscopy and image analysis were performed using the resources of the EPFL Bioimaging and Optics Core Facility (EPFL-BIOP). Mass spectrometry was performed at the Proteomics Core Facility (EPFL-PCF). We particularly thank Maria Pavlou and Mathilde Willemin for sharing their expertise in mass spectrometry.

## Author contributions

Conceptualization, MS, BM, DS; Methodology, MS, BM, DS; Software, BM; Validation, BM, JD, CD; Formal Analysis, MS, BM; Investigation, MS, BM, JD, CD; Data Curation, MS, BM; Writing — Original Draft, MS, BM, DS; Writing — Review & Editing, BM, JD, DS; Visualization, BM; Supervision, DS; Funding Acquisition, DS.

## Contact for reagents and resource sharing

Further information and requests for resource sharing and reagents should be directed to and will be fulfilled by the Lead Contact, David Suter.

## Data and Code availability

All original codes for reproducing analyses of this paper have been deposited on GitHub and are publicly available. Namely, key scripts for data analysis — inference algorithm and cell segmentation/fluorescence quantification, RNA-seq and dynamic SILAC analyses along with dynamic SILAC data — have been deposited on GitHub: https://github.com/UPSUTER/ProTuCo. LFQ proteomics were deposited on PRIDE under accession number PXD056601 (visible after login using login, password: VFrdb8hoSfb2). dSILAC proteomics data for NIH/3T3 cells were deposited on PRIDE under accession number PXD061384 (token: TDLxv360tAzh). dSILAC proteomics data for mESCs were deposited on PRIDE under accession number PXD061306 (token: m5mWMUMcgMTI). RNAseq data were deposited under accession number GSE278929 (token for reviewers: irsxcwqezdixhwv). Imaging data or other data reported in this paper are available from the lead contact upon request. Any additional information required to reanalyze the data reported in this paper is available from the lead contact upon request.

## STAR Methods

## EXPERIMENTAL MODEL AND SUBJECT DETAILS

### Cell Culture

NIH/3T3 and HEK293T cells were routinely cultured in DMEM (Gibco, 41966029), supplemented with 10% fetal bovine serum (Gibco, 10270-106), 1% penicillin/streptomycin (BioConcept, 401F00H) at 37°C, 5% CO_2_. Cells were passaged by trypsinization (Sigma, T4049-100ML) every 2-3 days and maintained at a confluency < 80%.

E14 and CGR8 mouse embryonic stem cell lines were routinely cultured in GMEM (Sigma-Aldrich, G5154) supplemented with 10% ES cell-qualified fetal bovine serum (Gibco, 16141– 079), 2 mM sodium pyruvate (Sigma-Aldrich, 113-24-6), 1% non-essential amino acids (Gibco, 11140–050), 1% penicillin/streptomycin (BioConcept, 4-01F00H), 2 mM L-glutamine (Gibco, 25030–024), 100 μM 2-mercaptoethanol (Sigma-Aldrich, 63689), leukaemia inhibitory factor (LIF — in-house), 3 μM GSK-3 Inhibitor XVI (Merck, 361559) and 0.8 μM PD184352 (Sigma-Aldrich, PZ0181), called hereafter 2i, at 37°C, 5% CO_2_. Cells were plated on dishes coated with gelatin (Sigma-Aldrich, G9391). Cells were passaged every 2-3 days by trypsinization (Sigma-Aldrich, T4049) when density reached approximately 3.0×10^4^ cells/cm^2^ (Mulas et al. 2019).

H1 human embryonic stem cells (WiCell Research Institute, WA01, 46XY) were routinely cultured as colonies in mTeSR Plus (STEMCELL Technologies, 100-0276) with 0.2% penicillin/streptomycin (BioConcept, 401F00H) at 37°C, 5% CO_2_. Cells were plated on dishes coated with Corning Matrigel hESC-qualified matrix (Corning, 354277). Cells were passaged every 4-7 days using the enzyme-free passaging reagent ReLeSR (STEMCELL Technologies, 05872) which preserves colonies, when most of the colonies were large with a dense center. For experiments requiring single cell dissociation, i.e. nucleofection, sorting, imaging, hESCs were passaged using Accutase (Innovative Cell Technology, AT104).

## METHOD DETAILS

### Production of lentiviral vectors and generation of stable cell lines

HEK293T cells were seeded at a density of 45,000 cells/cm^2^ and transfected using calcium phosphate one day after seeding. Cells were co-transfected with PAX2 (envelope), MD2G (packaging), and the lentiviral construct of interest, and concentrated 120-fold by ultracentrifugation as described previously^72^. Target NIH/3T3 cells were seeded at a density of 13,000/cm^2^ in a 24-well plate and transduced with 50 µL of concentrated lentivirus. For all dox-inducible cell lines, cells were first transduced with the pLV-pGK-rtTA3G-IRES-Bsd construct^73^, selected using 8 µg/mL of blasticidin (Thermofisher, A11139-03), transduced with the pLVTRE3G-construct of interest, and selected with puromycin (Thermofisher, A11138-03) at 2 µg/mL.

### Generation of the mESC SLT knock-in cell line

The SLT construct under the control of the EF1α promoter was first inserted using InFusion cloning (Takara) into the pDONOR MCS Rosa26 plasmid (Addgene #37200) containing Rosa26 homology harms. CGR8 cells were co-transfected with pDONOR-SLT, pCMV-RosaL6 ELD (Addgene #37198), and pCMV-RosaR4 KKR (Addgene #37199) with the lipofectamine 3000 transfection reagent (Invitrogen, L3000001) in a 6 well-plate following supplier instructions. These plasmids were a gift from Charles Gersbach^53^. Cells were then maintained for 2 weeks in culture. Cells were afterward sorted as single cells in 96-well plates for bright sfGFP and mOrange2 signals using a FACSAria (BD Biosciences) cell sorter. Around 1% of cells were positive. After 10 days of culture, single colonies were visually checked for MCFT fluorescence signal, and the brightest colonies were picked and amplified. 6 clones were then selected for characterization: gDNA was extracted using the DirectPCR Lysis Reagent (Viagen, 301-C) and PCRs were performed to verify the correct integration site of the transgene^74^ (Supplementary Table 4). After gel validation, the bands were cut, the DNA amplicons were extracted with QIAquick Gel Extraction Kit (QIAgene, 28704) and sent to Sanger sequencing (Microsynth) to check the integration site for the presence of indels.

### Generation of the hESC SLT knock-in cell line

The SLT construct was inserted using InFusion cloning (Takara) into the pC13N-dCas9-BFP-KRAB plasmid (Addgene, #127968), containing CLYBL homology arms and a CAG promoter. hESCs were co-transfected with pZT-C13-L1 (Addgene, #62196) and pZT-C13-R1 (Addgene, #62197) by nucleofection using the 4D-Nucleofector X Unit (Lonza) and Amaxa P3 Primary Cell kit L (Lonza, V4XP-3012), program CB-150. hESCs were supplemented with 1x CloneR2 (STEMCELL Technologies, 100-0691) supplement for the first two days. These plasmids were a gift from Jizhong Zou^45^. Cells were then maintained for 2 weeks in culture. Cells were afterward sorted as single cells in 96-well plates for bright sfGFP and mOrange2 signals using a FACS Aria (BD Biosciences) cell sorter. Around 0.4% of cells were positive. Cells were supplemented with 1x CloneR2 supplement for the first 4 days, with medium change every other day. After 14 days of culture, single colonies were visually checked for SLT signal and the brightest colonies were picked and amplified. 3 clones were then selected for characterization: gDNA was extracted using the DNEasy Blood & Tissue kit (Qiagene, 69504) and PCRs were performed to verify the correct integration site of the transgene (Supplementary Table 4). After running PCR products on an agarose gel, the bands were cut, the DNA amplicons were extracted using QIAquick Gel Extraction Kit (Qiagene, 28704) and verified by Sanger sequencing (Microsynth).

### Differentiation of hESCs into astrocytes

hESCs were first differentiated to neural progenitor cells using the STEMdiff SMADi Neural Induction kit (STEMCELL Technologies, 08581) according to the manufacturer’s protocol for monolayer culture. Briefly, 2 x 10^5^ cells/cm^2^ were plated as single cells on Corning Matrigel hESC-qualified matrix (Corning, 354277) in STEMdiff Neural Induction Medium with SMADi supplement and 10 µM ROCK Inhibitor (Y-27632) (MilliporeSigma, SCM075). The next day, Y-27632 was removed, and the medium was changed daily. Cells were passaged three times every 7 days using Accutase (Innovative Cell Technology, AT104). One day after the third passage, the medium was changed to STEMdiff Astrocyte Differentiation Medium kit (STEMCELL Technologies, 100-0013) to obtain astrocyte precursors. The astrocyte precursors were maintained for three additional passages, changing the medium daily during the first week and every 2 – 3 days for the remaining weeks. Following three weeks differentiation after the third passage of the precursors, the STEMdiff Astrocyte Maturation medium was used (STEMCELL Technologies, 100-0016). Maturation of astrocytes was performed for another three weeks, performing single cell passage every 7 days and medium change every 2 – 3 days, before the downstream analysis.

### Immunofluorescence for characterization of astrocyte-enriched cell culture

Cells were first fixed using 4 % formaldehyde (FA) (Thermo Fischer Scientific, 28906) in PBS that was added 1:1 to the culture medium and incubated for 5 min at room temperature (final FA: 2%). The medium was removed, and 4 % FA was added for 15 min at room temperature. After two washes with PBS, cells were permeabilized with 0.1 % Triton X-100 (BioChemica, UN3082) in PBS for 20 min and blocked with 1 % BSA (Sigma-Aldrich, A7906) in PBS for 30 min. Cells were incubated overnight at 4 °C with anti-Ki67 antibody (1:100, BD Biosciences, 550609) and anti-GFAP antibody (1:200, STEMCELL Technologies, 60048). The next day, cells were washed twice with PBS and incubated with an anti-mouse secondary antibody conjugated to AlexaFluor647 (1:1000, LifeTechnologies, A31571) for 1 h at room temperature. Cells were washed twice with PBS and mounted with Vectashield-containing DAPI (Vector, H-1500-10). Imaging was performed with an Operetta CLS (Perkin Elmer) microscope, 20× objectives (Air immersion, Plan Apochromat, NA 0.8), at RT, using the following filters: Ex: BP 615-645, BP 650-675, Em: HC 655-760 for the antibody channels.

### Snapshot imaging of the MCFT upon CHX/MYCi/INK128 steady-state treatments

3 days before imaging, NIH/3T3 cells were seeded on a 96-well plate (Perkin Elmer, 6055302) coated with fibronectin (Sigma, F4759) at a density that does not exceed 80% confluency for each well when imaging starts. After seeding, cells were cultured overnight without CHX in FluoroBrite™ DMEM medium (Gibco, A1896701) supplemented with 10% fetal bovine serum (Gibco, 10270-106), 2 mM L-glutamine (Gibco, 25030–024), and 1% penicillin/streptomycin (BioConcept, 401F00H). The next day, CHX treatment was applied to each well following the designed concentration gradients (Supplementary Table 1). The CHX concentration gradient was made of 56 different concentrations ranging from 0.002 to 0.5 µg/mL. The medium containing the respective CHX concentrations was changed once after 24 hours. After 2 days of treatment, snapshot imaging was performed with Operetta CLS (Perkin Elmer), 20× objectives (Air immersion, Plan Apochromat, NA 0.8), at 37 °C, 5% CO_2_. For the sfGFP channel, the following filters were used: Ex: BP 435-460, 460-490, Em: HC 500-550. For the mOrange2 channel, the following filters were used: Ex: BP 490-515, 530-560, Em: HC 570-650.

Experiments with H1 hESCs, astrocytes and CGR8 mESCs were performed similarly, except for culture conditions. hESCs were cultured in mTeSR1 without phenol red (STEMCELL Technologies, 05876) and seeded on a 96-well plate coated with Corning Matrigel hESC-qualified matrix (Corning, 354277). A 10 µM ROCK Inhibitor (Y-27632) (MilliporeSigma, SCM075) was added at the time of seeding. Mature astrocytes were seeded on a 96-well plate coated with Corning Matrigel hESC-qualified matrix (Corning, 354277) in Brain-Phys Without Phenol-Red (STEMCELL Technologies, 05791) supplemented with STEMdiff Astrocyte Maturation Supplement A and B (STEMCELL Technologies, 100-0037 and 100-0017). mESCs were cultured in phenol-red free N2B27 + 2i/LIF medium and seeded on a 96-well plate coated with laminin 511 (BioLamina, LN511-0202). Before starting CHX treatment, mESCs were cultured for at least 2 weeks in N2B27 + 2i/LIF medium.

A similar protocol was used for anisomycin and MYCi treatments. The anisomycin concentration gradient was made of 56 different concentrations ranging from 0.0002 to 0.05 µg/mL (Supplementary Table 2). The MYCi concentration gradient was made of 56 different concentrations ranging from 0.26 to 64 µM (Supplementary Table 3).

For CHX treatment in combination with INK128, we proceeded as previously described, adding a constant concentration of INK128 to the CHX plate gradient.

### Live imaging of the MCFT upon CHX treatment/release

3 days before imaging, NIH/3T3 cells were seeded on a 96-well plate coated with fibronectin (Sigma, F4759) at a density that does not exceed 80% confluency for each well when imaging starts. Four conditions were designed: control (-CHX), CHX (2 days treatment with 0.1 µg/ml CHX before imaging), pulse (0.1 µg/ml CHX added right before imaging), and release (2 days of treatment with 0.1 µg/ml CHX before imaging followed by CHX removal immediately before imaging). 500 cells were seeded per well for control and pulse conditions. 1000 cells were seeded per well for CHX and release conditions. After seeding, cells were cultured overnight without CHX in FluoroBrite™ DMEM (Gibco, A1896701) supplemented with 10% fetal bovine serum (Gibco, 10270-106), 2 mM L-glutamine (Gibco, 25030–024), and 1% penicillin/streptomycin (BioConcept, 401F00H). The next day, CHX was applied at 0.1 µg/ml in CHX and release conditions. During the treatment, the medium was changed every 24h. Before imaging, all wells were washed once with medium. Then, medium containing 0.1 µg/ml CHX was added to CHX and pulse conditions. In control and release conditions, medium without CHX was added. Live imaging was performed with Operetta CLS (Perkin Elmer), 20× objectives (Air immersion, Plan Apochromat, NA 0.8), at 37 °C, 5% CO_2_, with 15 min intervals for more than 20 h if not specified otherwise. For the sfGFP channel, the following filters were used: Ex: BP 435-460, 460-490, Em: HC 500-550. For the mOrange2 channel, the following filters were used: Ex: BP 490-515, 530-560, Em: HC 570-650.

Experiments with H1 hESCs, human astrocytes, and CGR8 mESCs were performed similarly, except for the culture conditions that were as described in the previous section.

For the CHX pulse experiment in the presence of 25 nM INK128 or 200 nM ISRIB, we proceeded as previously described, pre-treating the cells 1 h with the respective drugs before adding the medium with CHX, keeping constant the INK128/ISRIB concentration.

### Live imaging of the MCFT upon MYCi treatment/release

3 days before imaging, NIH/3T3 cells were seeded on a 96-well plate coated with fibronectin (Sigma, F4759) at a density that does not exceed 80% confluency for each well when imaging starts. Four conditions were designed: control (-MYCi), MYCi (2 days treatment with 64 µM MYCi before imaging), pulse (64 µM MYCi added right before imaging), and release (2 days of treatment with 64 µM MYCi before imaging followed by MYCi removal immediately before imaging). 254 cells were seeded per well for control and pulse conditions. 800 cells were seeded per well for MYCi and release conditions. After seeding, cells were cultured overnight without MYCi in FluoroBrite™ DMEM (Gibco, A1896701) supplemented with 10% fetal bovine serum (Gibco, 10270-106), 2 mM L-glutamine (Gibco, 25030–024), and 1% penicillin/streptomycin (BioConcept, 401F00H). The next day, MYCi was applied at 64 µM in MYCi and release conditions. During the treatment, the medium was changed every 24h. Before imaging, all wells were washed once with medium. Then, medium containing 64 µM MYCi was added to MYCi and pulse conditions. In control and release conditions, medium without MYCi was added. Live imaging was performed with Operetta CLS (Perkin Elmer), 20× objectives (Air immersion, Plan Apochromat, NA 0.8), at 37 °C, 5% CO_2_, with 15 min intervals for more than 20 h if not specified otherwise. For the sfGFP channel, the following filters were used: Ex: BP 435-460, 460-490, Em: HC 500-550. For the mOrange2 channel, the following filters were used: Ex: BP 490-515, 530-560, Em: HC 570-650.

### SNAP-tag pulse-chase labeling

Cells were seeded and cultured in 96-well plates following the previous live imaging method section. 45 minutes before imaging, cells were incubated with an imaging medium that contained 40 - 80 nM SNAP Cell 647 Sir dye (NEB, S9102S) for 30 min at 37°C, 5% CO_2_. Cells were then gently washed five times with fresh medium. The medium was then replaced with a fresh imaging medium containing 1 µM SNAP Cell Block (NEB, S9106S) to prevent the binding of residual SNAP dye to newly synthesized SNAP-tagged proteins. Imaging was performed with an Operetta CLS (Perkin Elmer) microscope, 20× objectives (Air immersion, Plan Apochromat, NA 0.8), at 37 °C 5% CO_2_, at 15 min intervals. For the sfGFP channel, the following filters were used: Ex: BP 435-460, 460-490, Em: HC 500-550. For the SNAP channel, the following filters were used: Ex: BP 615-645, BP 650-675, Em: HC 655-760.

### Dox pulse-chase experiment

Three days before imaging, NIH/3T3 cells were seeded at a density of 200 cells/well on a 96- well plate coated with fibronectin (Sigma, F4759). After seeding, cells were cultured overnight without dox in FluoroBrite™ DMEM (Gibco, A1896701) supplemented with 10% fetal bovine serum (Gibco, 10270-106), 2 mM L-glutamine (Gibco, 25030–024), and 1% penicillin/streptomycin (BioConcept, 401F00H). The next day, dox (Sigma-Aldrich, D9891) treatments were applied at the desired concentrations. During the treatment, the medium was changed every 24 hours. Before imaging, all wells were washed three times with warm PBS followed by the addition of medium containing the desired dox concentration. Live imaging was performed thereafter (∼15min delay) with an Operetta CLS (Perkin Elmer), 20× objectives (Air immersion, Plan Apochromat, NA 0.8), at 37 °C, 5% CO_2_, with 15 min intervals for at least 20 h if not specified otherwise. For the sfGFP channel, the following filters were used: Ex: BP 435-460, 460-490, Em: HC 500-550. For the mOrange2 channel, the following filters were used: Ex: BP 490-515, 530-560, Em: HC 570-650.

### Imbalance computation

The imbalance *θ*, introduced in ^50^ is defined as:

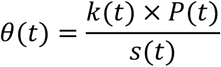

The protein synthesis and decay rates at time *t*, *s(t)*, and *k(t)* respectively, were inferred using the Hierarchical Bayesian inference algorithm introduced hereafter (see also Supplementary Text for details). The total protein level at time *t*, *P(t)*, was calculated by summing up the measured green *G* signal and the inferred black-green *B_G_* signal such that:

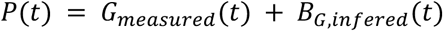

### *N*-Hydroxysuccinimide (NHS ester) labeling and imaging

Cells were seeded in a 96-well plate coated with fibronectin (Sigma, F4759) and cultured overnight. The cell seeding number was adjusted so that none of the wells would be more than 80% confluent on day 7. On day 2, 48 different CHX concentrations ranging from 0.002 to 0.25 µg/mL were applied to cells for the next 6 days. The medium was changed every 48 hours. On day 7, cells were fixed with 4% formaldehyde in PBS for 15 min at RT before washing once with PBS. Then, cells were permeabilized with 100% pre-cooled methanol at -20°C for 10 min. Before staining, cells were washed once with 0.2 M sodium bicarbonate. N-Hydroxysuccinimide (Invitrogen, A37573) was diluted in 0.2 M sodium bicarbonate to a final concentration of 50 µg/mL and applied to cells at RT for 30 min. Cells were washed twice with PBS and mounted with Vectashield containing DAPI (Vector, H-1500-10). Imaging was performed with an Operetta CLS (Perkin Elmer) microscope, 20× objectives (Air immersion, Plan Apochromat, NA 0.8), at RT, using the following filters: Ex: BP 615-645, BP 650-675, Em: HC 655-760 for the NHS-ester channel.

### ELISA for p70S6K

To measure phosphorylation levels of p70 S6K in NIH/3T3 and mES cells, we used the Multispecies p70 S6 Kinase (Total/Phospho) InstantOne™ ELISA Kit (Invitrogen). We followed the manufacturer’s assay protocol for adherent cells. Briefly, we plated 5,000 to 10,000 (48 h timepoint) cells per well of a 96-well plate and treated the cells with 0.1 µg/mL CHX for different durations (0, 1, 2, 5 48 hours). After 48 hours, we performed cell lysis using 150 µL per well of the Cell Lysis Buffer Mix (1X) supplemented with benzonase. We then proceeded as described by the manufacturer.

### RNA-seq

#### Sample collection

NIH/3T3 and mES cells were cultured in the standard conditions described above. At timepoint 0 h, CHX at a final concentration of 0.1µg/mL was pulsed into the medium. At subsequent timepoints (1 h, 2 h, 5 h, 48 h), cells were washed once in PBS and directly lysed using the lysis buffer of the RNeasy Mini kit (Quiagen). RNA extraction was then performed according to the manufacturer instructions. RNA quality and concentration was assessed using Nanodrop and TapeStation 4200 (Agilent), which confirmed their integrity.

#### Sample preparation and sequencing

Libraries for mRNA-seq were prepared with the Stranded mRNA Ligation method (Illumina), according to manufacturer’s instructions, starting from 300ng RNA. Libraries, all bearing unique dual indexes, were subsequently loaded at 9.9 pM on an Aviti Cloudbreak Freestyle flow cell (Element Biosciences) and sequenced according to manufacturer instructions, yielding pairs of 80 nucleotides reads at a depth of about 30 mio reads pairs per sample. Reads were trimmed of their adapters with bases2fastq version 1.8.0.1260801529 (Element Biosciences) and quality-controlled with fastQC v0.11.9. FastQ Screen v0.14.0 tool was used for screening FASTQ files reads against multiple reference genomes.

#### Data analysis

Indexes were generated with STAR^75^ using GRCm39 mouse genome assembly with option -- sjdbOverhang 100. The reads were then aligned using STAR using the --quantMode GeneCounts option. A count matrix was then generated using an in-house Python script that takes as input the ReadsPerGene.out.tab files generated by STAR alignment. DESeq2^76^ was then used in R 4.4.1^77^ in order to perform differential expression analysis. The output was then analyzed in Python using standard libraries. Lists of ribosomal and mitochondrial genes were respectively taken from ^78^ and ^79^.

### Label-free Mass Spectrometry experiments

#### Sample collection

In each condition (DMSO or 0.1 µg/mL CHX), one 10 cm culture dish at <80% confluency was used. Cells were washed twice in PBS without Mg^2+^ and Ca^2+^ (PBS -/- hereafter), trypsinized, collected in a 15 mL tube, and pelleted by centrifugation at 300g for 5 min. The supernatant was then discarded, and the cell pellet was snap-freezed in liquid nitrogen. Once all samples were collected, the pellets were resuspended in lysis buffer (100 mM Tris buffer pH 8, 2% SDS, 1X Halt^TM^ Protease Inhibitor and 1:50 v:v Benzonase nuclease) and incubated 15 min at room temperature. The samples were then boiled at 90°C for 10 min and centrifuged at 16’000g for 10 min at 4°C. The supernatant was then transferred to a new tube and the protein extract concentration measured using a BCA assay (Pierce).

#### Sample preparation for Mass Spectrometry

Mass spectrometry-based proteomics-related experiments were performed by the Proteomics Core Facility at EPFL. Each sample was digested by filter aided sample preparation (FASP)^80^ with minor modifications. Proteins (20 µg) were deposited on top of washed and conditioned Microcon®-30K devices (Merck AG, Zug, Switzerland). Samples were centrifuged at 9400×*g*, at 20°C for 30 min. or until complete dryness. All subsequent centrifugation steps were performed using the same conditions. Two washing steps were performed using 200 μL urea solution (8 M Urea, 100 mM Tris-HCl pH 8). Reduction was performed by adding 100 μL of 10 mM Tris(2-carboxy)phosphine (TCEP) in urea solution on top of filters followed by 60 min incubation time at 37°C with gentle shaking and light protection. Reduction solution was removed by centrifugation and two washing steps with 200 μL urea solution. Then, alkylation was performed by adding 100 μL of 40 mM chloroacetamide (CAA) in urea solution and incubating the filters at 37°C for 45 min with gentle shaking and protection from light. The alkylation solution was removed by centrifugation followed by two washing steps with 200 μL of urea solution. Finally, two additional washing steps using 200 μL of 5 mM Tris-HCl pH 8 were performed to condition the filters for digestion. Proteolytic digestion was performed overnight at 37°C by adding 100 μL of a combined solution of Endoproteinase Lys-C and Trypsin Gold in an enzyme/protein ratio of 1:50 (w/w) prepared in 5 mM Tris-HCl and 10 mM CaCl_2_ on top of filters. The resulting peptides were recovered by centrifugation and two subsequent elution with 50 μL of 4% trifluoroacetic acid. Finally, the recovered peptides were desalted on SDB-RPS StageTips^81^ and dried by vacuum centrifugation prior to LC-MS/MS injections.

#### Mass spectrometry

Samples were resuspended in 2% acetonitrile (Biosolve), 0.1% FA and nano-flow separations were performed on a Dionex Ultimate 3000 RSLC nano UPLC system (Thermo Fischer Scientific) on-line connected with an Exploris Orbitrap 480 Mass Spectrometer (Thermo Fischer Scientific). A capillary precolumn (Acclaim Pepmap C18, 3 μm-100Å, 2 cm x 75μm ID) was used for sample trapping and cleaning. A 50 cm long capillary column (75 μm ID; in-house packed using ReproSil-Pur C18-AQ 1.9 μm silica beads; Dr. Maisch) was then used for analytical separations at 250 nl/min over 150 min biphasic gradients. Acquisitions were performed through Top Speed Data-Dependent acquisition mode using a cycle time of 2 seconds. First MS scans were acquired with a resolution of 60’000 (at 200 m/z) and the most intense parent ions were selected and fragmented by High energy Collision Dissociation (HCD) with a Normalized Collision Energy (NCE) of 30% using an isolation window of 1.4 m/z. Fragmented ions were acquired with a resolution 15’000 (at 200m/z) and selected ions were then excluded for the following 20 s.

#### Data analysis

Raw data were processed using MaxQuant 1.6.10.43^82^ against 55286 entries (LR2022_05), Carbamidomethylation was set as fixed modification, whereas oxidation (M), phosphorylation (S, T, Y), acetylation (Protein N-term) and glutamine to pyroglutamate were considered as variable modifications. A maximum of two missed cleavages were allowed and “Match between runs” option was enabled. A minimum of 2 peptides was required for protein identification and the false discovery rate (FDR) cutoff was set to 0.01 for both peptides and proteins. Label-free quantification and normalization was performed by MaxQuant using the MaxLFQ algorithm, with the standard settings^83^. Statistical analysis was performed using Perseus version 1.6.12.0^84^ from the MaxQuant tool suite. Reverse proteins, potential contaminants, and proteins only identified by sites were filtered out. Protein groups containing at least 3 valid values in at least one condition were conserved for further analysis. Empty values were imputed with random numbers from a normal distribution (Width: 0.4 and Down shift: 1.8 std). A two-sample t-test with permutation-based FDR statistics (250 permutations, FDR = 0.01, S0 = 1) was performed to determine significant differentially abundant candidates. Lists of ribosomal and mitochondrial proteins were respectively taken from ^78^ and ^79^.

### Dynamic SILAC

#### Cell culture

NIH/3T3 cells were cultured at least 2 weeks in light medium: DMEM for SILAC (Thermo scientific) supplemented with PS, dialyzed FBS (Thermo scientific), (light) L-Lysine-2HCl (0.666 mM), (light) L-Arginine-HCl (0.399 mM)^44^, and L-Proline (200 mg/L)^85^. mESCs were cultured at least 2 weeks in medium-heavy medium: N2B27 for SILAC (DMEM/F12 for SILAC (AthenaES), Neurobasal for SILAC (AthenaES), Sodium Pyruvate (40 mg/mL), N2 (1X), B27 (0.5X), Pen/Strep (1%), L-glutamine (2 mM), beta-mercaptoethanol (50 µM)) + 2i/LIF supplemented with (medium-heavy) ^13^C_6_ ^15^N_4_ L-arginine (0.65 mM), (medium-heavy) ^13^C_6_ L-arginine (0.55 mM), and L-Proline (200 mg/L)^85^. Before switching from light (NIH/3T3) or medium-heavy (mESC) to heavy medium, cells were treated with 0.05 µg/mL CHX or equivalent dilution of DMSO in 6-well plates. Note that we decided to treat the cells with 0.05 µg/mL CHX instead of the usual 0.1 µg/mL concentration due to the lower division rate of NIH/3T3 in the SILAC medium. At timepoint 0 h, we replaced light or medium-heavy medium with pre-warmed heavy medium: DMEM for SILAC (Thermo scientific) supplemented with PS, dialyzed FBS (Thermo scientific), (heavy) ^13^C_6_ ^15^N_2_ L-Lysine-2HCl (0.666 mM), (heavy) ^13^C_6_ ^15^N_4_ L-Arginine-HCl (0.399 mM), and L-Proline (200 mg/L) for NIH/3T3 ; N2B27 for SILAC (Thermo scientific) supplemented with (heavy) ^13^C_6_ ^15^N_2_ L-Lysine-2HCl (0.65 mM), (heavy) ^13^C_6_ ^15^N_4_ L-Arginine-HCl (0.55 mM), and L-Proline (200 mg/L) for mESC.

#### Sample collection

At the corresponding timepoint, the culture dish was placed on ice, the medium aspirated, and the cells washed twice with ice-cold PBS without Mg^2+^ and Ca^2+^ (PBS -/- hereafter). Cells were then scraped in 50 µL of lysis buffer (100 mM Tris buffer pH 8, 2% SDS, 1X Halt^TM^ Protease Inhibitor and 1:50 v:v Benzonase nuclease), collected in a 1.5 mL Eppendorf *protein LoBind* tube, and incubated 15 min at room temperature. The protein extract was then snap-freezed in liquid nitrogen. Once all samples were collected, the samples were boiled at 90°C for 10 min and centrifuged at 16,000xg for 10 min at 4°C. The supernatant was then transferred to a new tube, and the protein extract concentration was measured using a BCA assay (Pierce).

#### Sample preparation for Mass Spectrometry

Protein samples were prepared as described above for the Label-free Mass Spectrometry experiments.

#### Mass spectrometry

Samples were resuspended as defined earlier but the separation was performed using a Vanquish Neo nano UPLC system (Thermo Fischer Scientific) on-line connected with an Orbitrap Fusion Lumos Tribrid Mass Spectrometer (Thermo Fischer Scientific). The same parameters were used for the separation on the gradient but the ones of the MS acquisition were slightly different. Acquisitions were performed through Top Speed Data-Dependent acquisition mode using a cycle time of 1 second. First MS scans were acquired with a resolution of 240K (at 200 m/z) on the orbitrap and the most intense parent ions were selected and fragmented by High energy Collision Dissociation (HCD) with a Normalized Collision Energy (NCE) of 30% using an isolation window of 0.7 m/z. Fragmented ions were acquired on the ion trap with a maximum injection of 20 ms and selected ions were then excluded for the following 20 s.

#### Data analysis

Raw data were processed using MaxQuant 2.4.4.0^82^ against 54822 entries (LR2024_01) using the parameters taken from ^64^ with slight modifications as described in Tables a and b below:

**Table a:** MaxQuant parameters used for dSILAC in NIH/3T3 cells.

| Category | Parameter | A: Turnover analysis (dynamic SILAC) | B: (Re-)incorporation of light amino acids | C: LFQ analysis of t=0 samples |
| --- | --- | --- | --- | --- |
| **Group specific parameters** |  |  |  |  |
| **Type** | Type | Standard | Standard | Standard |
|  | Multiplicity | 2 | 1 | 1 |
|  | Max labeled AAs | 3 | - | - |
|  | Labels | Light: -<br>Heavy: Arg10, Lys8 | - | - |
| **Modifications** | Variable | Oxidation (M)<br>Acetyl (N-term Protein) | Oxidation (M)<br>Acetyl (N-term Protein)<br>Arg10 as PTM<br>Lys8 as PTM | Oxidation (M)<br>Acetyl (N-term Protein)<br>Gln -> pyro-Glu<br>Phospho (STY) |
|  | Fixed | Carbamidomethyl (C) | Carbamidomethyl (C) | Carbamidomethyl (C) |
|  | Max nb of modif. | 5 | 5 | 5 |
| **Label-free quantification** | LFQ | None | None | LFQ |
|  | LFQ min. ratio count | - | - | 2 |
|  | Normalisation type | - | - | Classic |
| **Misc.** | Requantify | FALSE | FALSE | FALSE |
| **Digestion** | Digestion mode | Specific | Specific | Specific |
|  | Enzyme | Trypsin | Trypsin | Trypsin |
|  | Max. missed cleavages | 2 | 2 | 2 |
| **Global parameters** |  |  |  |  |
| **Sequences** | Fasta files | Uniprot_Mouse_54822Sequences_LR2024_01 |  |  |
|  | Include contaminants | TRUE | TRUE | TRUE |
| **Protein quantification** | Label min. ratio count | 2 | 1 | 2 |
|  | Peptides for quantification | Unique + razor | All | Unique + razor |
| **Identification** | Min. peptides | 2 | 1 | 2 |
|  | Min. Razor + unique peptides | 2 | 1 | 2 |
|  | Min. unique peptides | 1 | 0 | 1 |
|  | Match between runs | TRUE | TRUE | TRUE |

**Table b:** MaxQuant parameters used for dSILAC in mESC.

Table a: MaxQuant parameters used for dSILAC in NIH/3T3 cells.
| Category | Parameter | A: Turnover analysis (dynamic SILAC) | B: (Re-)incorporation of light amino acids | C: LFQ analysis of t=0 samples |
| --- | --- | --- | --- | --- |
| **Group specific parameters** |  |  |  |  |
| **Type** | Type | Standard | Standard | Standard |
|  | Multiplicity | 2 | 1 | 1 |
|  | Max labeled AAs | 3 | - | - |
|  | Labels | Light: Arg6, Lys4<br>Heavy: Arg10, Lys8 | - | Arg6<br>Lys4 |
| **Modifications** | Variable | Oxidation (M)<br>Acetyl (N-term Protein) | Oxidation (M)<br>Acetyl (N-term Protein)<br>Arg6 as PTM<br>Arg10 as PTM<br>Lys4 as PTM<br>Lys8 as PTM | Oxidation (M)<br>Acetyl (N-term Protein)<br>Gln -> pyro-Glu<br>Phospho (STY) |
|  | Fixed | Carbamidomethyl (C) | Carbamidomethyl (C) | Carbamidomethyl (C) |
|  | Max nb of modif. | 5 | 5 | 5 |
| **Label-free quantification** | LFQ | None | None | LFQ |
|  | LFQ min. ratio count | - | - | 2 |
|  | Normalisation type | - | - | Classic |
| **Misc.** | Requantify | FALSE | FALSE | FALSE |
| **Digestion** | Digestion mode | Specific | Specific | Specific |
|  | Enzyme | Trypsin | Trypsin | Trypsin |
|  | Max. missed cleavages | 2 | 2 | 2 |
| **Global parameters** |  |  |  |  |
| **Sequences** | Fasta files | Uniprot_Mouse_54822Sequences_LR2024_01 |  |  |
|  | Include contaminants | TRUE | TRUE | TRUE |
| **Protein quantification** | Label min. ratio count | 2 | 1 | 2 |
|  | Peptides for quantification | Unique + razor | All | Unique + razor |
| **Identification** | Min. peptides | 2 | 1 | 2 |
|  | Min. Razor + unique peptides | 2 | 1 | 2 |
|  | Min. unique peptides | 1 | 0 | 1 |
|  | Match between runs | TRUE | TRUE | TRUE |

#### Computation of protein half-life

We provide a detailed description of the analysis steps in the Supplementary Text and the scripts deposited on GitHub. Briefly, we followed ^64^ by first filtering out contaminants, decoy sequences, and peptides exhibiting false heavy signal in the unlabeled control sample. We then computed the percentage of old (light) peptides *%old* according to:

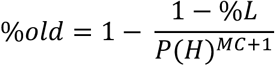

In which *%L* is the fraction of remaining light peptide, *P(H)* is the probability of heavy Lys/Arg incorporation, and *MC* is the number of missed cleavages. Considering *P(H)*, we correct for light amino acid recycling. In the experiment presented in the main text for NIH/3T3, we assumed *P(H)∼1* since the measured *P(H)=0.985*. For mESC, *P(H) = 0.95*. Grouping peptides per protein group, we then performed a linear fit according to the following equation:

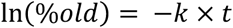

In which *k* is the decay rate and *t* is the incubation time in the heavy medium. We kept protein groups for which *r^2^>0.9* following ^44^. Finally, we computed the protein half-life using the following equation:

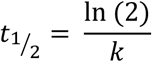

### Image pre-processing

For background correction, image intensity was modeled the following way:

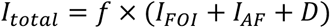

*I_total_* is the intensity value of the raw image. *f* is the uneven illumination pattern. *I_FOI_* is the fluorescence signal of interest. *I_AF_* is the auto-fluorescence from the medium. *D* is the dark field signal. *D* was ignored as it was negligible compared to *I_FOI_* or *I_AF_*. *f* was generated by imaging a well with medium only using the same exposure settings. This image was subsequently normalized to its mean pixel intensity. Raw images were divided by the image from the well with medium only to generate flat-field corrected images. *I_AF_* was calculated for every single frame by applying the appropriate thresholding method to the field-corrected images, which created a binary image that masked foreground signals *I_FOI_*. The threshold was determined using the fifth percentile of the intensity distribution of the pixels belonging to segmented cells. The mask was then enlarged by erosion. Finally, the mean or the peak of pixel intensities was measured from the unmasked region. We did not generate background-subtracted images but rather subtracted this value from final single-cell measurements inside a Jupyter notebook.

### Cell segmentation and tracking

For segmentation and tracking nuclear fluorescent signals, Trackmate (7.1)^86^ was used through a groovy script (written by Olivier Burri and Romain Guiet, EPFL) that enables image processing in batch. This script was tailored for using Stardist^87^ as the detector to segment cell nuclei on the sfGFP channel. Tracking was then performed with the LAP algorithm^88^ within the same script. The script generated Trackmate XML files that allow to review the tracking result with the Trackmate user interface. Both sfGFP and mOrange2 traces were smoothed with a Savitzky-Golay filter (window length: 100, polynomial order: 6)^89^ before superstatistical modeling. For the CHX pulse and release experiment in hESCs and mESCs, fluorescence trajectories were detrended using the Control and the +CHX conditions, respectively. For the NHS-ester labeling and mESC/hESC experiments, CellPose in Python^90^ was used to segment the nuclei. The masks were then used to retrieve NHS-ester SiR-647 nm integrated intensities of individual cells. All microscopy figure panels and supplementary videos were made using the microfilm package^91^ in Python.

### Quantification of *k* and *k_deg_* from SNAP pulse-chase labeling

To measure *k_deg_* with SNAP, the integrated intensity for each single cell trajectory or single lineage (mother and computationally-fused daughter cells) trajectory was transformed by natural logarithm, then linear-fitted a robust RANSAC^92^ regression (random state: 42) from the scikit-learn package 1.0.2^93^. Fluorescence intensities of daughter cells resulting from cell division were summed. To measure the total protein decay rate *k* with SNAP (*k_deg_* + *k_dil_)*, the mean intensities for each lineage were transformed by natural logarithm and then linear fitted with a robust regressor RANSAC (random state: 42). Fluorescence intensities of daughter cells resulting from cell division were averaged.

### Quantification of *S* and *k* from the MCFT

The decay rate *k* was computed using the formula derived in the Supplementary Text. The procedure to calibrate the MCFT is given in the Supplementary Text. To compute the rates *s*, we assumed equilibrium for all our measurements. With *μ_G_*, *μ_R_*, the integrated intensity for sfGFP and mOrange2 fluorescence, and *m_G_* the maturation rate of sfGFP (Fig.1A), we computed the synthesis rate as:

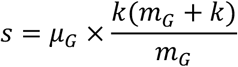

*s* is thus given here in relative units (fluorescence/hour). In the main text, we use *S* when *s* is normalized to the control condition, such that for control conditions we have *S=1*. See Supplementary Text for details. To compute *s_conc_*. we used the same equation, replacing the integrated intensity *μ_G_* with the mean intensity 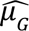 (average fluorescence intensity per pixel).

The equation linking the two observables is:

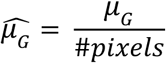

In the main text, we use *S_conc._* when *s*_;4+;._ is normalized to the median synthesis rate in a cell line (Fig.5).

### Quantification of *k_dil_* from time-lapse movies

To measure *k_dil_*, we counted cell numbers per imaging frame over time. The dilution rate was then calculated by log2 transformation of the cell number and linear fitting with the RANSAC regressor (random state: 42).

### Modeling and inference from MCFT traces

We adopted a superstatistical Bayesian inference algorithm^47–49^, designed for autoregressive models (AR-1), and applied it to our ordinary differential equation (ODE) systems. Briefly, we computed the (joint) posterior distribution of *S* and *k* along with the latent variable *B_G_* at each timepoint and propagated this posterior forward and backward along the MCFT trajectories. In our case, the trajectories used for the inference are *G* (sfGFP) and *R* (mOrange2) fluorescence trajectories acquired by live-cell imaging. Propagation of the posterior distribution relies on two hyperparameters chosen for their ability to recapitulate the data (data retrodiction) and fixed for a whole dataset. Details are given in the Supplementary text.

### Log-likelihood and bootstrapping for model selection

Log-likelihood was computed using a Gaussian error model. The pseudo-(log) likelihood was used for the actual computation, finally reducing to the residual sum of squares. Briefly, data were compared to the model predictions for the passive adaptation, no-adaptation, and perfect adaptation models. To compute statistics for the log-likelihood estimator, bootstrapping was performed with N=1000 resampling, keeping the sample size constant.

## STATISTICAL ANALYSIS

Statistical analysis was performed using Python and the scipy.stats package. All details about statistical tests and sample size are available in the figures’ legends. p<0.05 was considered statistically significant. When Pearson correlation was performed, the statistics (p-value and determination coefficient *r^2^*) were computed and displayed on the figure or specified in the figure legend. Data binning was performed using an in-house algorithm. Briefly, the number of bins was fixed over the full range of values of the x-axis. For each bin, we then computed the median of the y-values of all data points belonging to it.

### Ethics statement

All experiments involving hESC were approved by the Canton of Vaud Ethics committee on human research (https://www.cer-vd.ch).

## References

1. Dikic, I. & Elazar, Z. Mechanism and medical implications of mammalian autophagy. Nat. Rev. Mol. Cell Biol. 19, 349–364 (2018).

2. Balakrishnan, R. et al. Principles of gene regulation quantitatively connect DNA to RNA and proteins in bacteria. Science 378, eabk2066 (2022).

3. Seinkmane, E. et al. Circadian regulation of protein turnover and proteome renewal. 2022.09.30.509905 Preprint at 10.1101/2022.09.30.509905 (2022).

4. Klein, M. E., Castillo, P. E. & Jordan, B. A. Coordination between Translation and Degradation Regulates Inducibility of mGluR-LTD. Cell Rep. 10, 1459–1466 (2015).

5. Ramachandran, K. V. et al. Activity-Dependent Degradation of the Nascentome by the Neuronal Membrane Proteasome. Mol. Cell 71, 169–177.e6 (2018).

6. Kristensen, A. R., Gsponer, J. & Foster, L. J. Protein synthesis rate is the predominant regulator of protein expression during differentiation. Mol. Syst. Biol. 9, 689 (2013).

7. Honda, T., et al. Coordination of gene expression with cell size enables Escherichia coli to efficiently maintain motility across conditions. Proc. Natl. Acad. Sci. 119, e2110342119 (2022).

8. Liu, S. et al. Large cells activate global protein degradation to maintain cell size homeostasis. 2021.11.09.467936 Preprint at 10.1101/2021.11.09.467936 (2021).

9. Motil, K. J. et al. Whole-body leucine and lysine metabolism: response to dietary protein intake in young men. Am. J. Physiol. 240, E712–721 (1981).

10. Wu, C.-I. & Wen, H. Heightened protein-translation activities in mammalian cells and the disease/treatment implications. Natl. Sci. Rev. 7, 1851–1855 (2020).

11. Wolfe, R. R. Effects of amino acid intake on anabolic processes. Can. J. Appl. Physiol. Rev. Can. Physiol. Appl. 26 **Suppl**, S220–227 (2001).

12. Tg, G., Ms, B., Pt, R., E, V. & Bb, R. Essential amino acid ingestion alters expression of genes associated with amino acid sensing, transport, and mTORC1 regulation in human skeletal muscle. Nutr. Metab. 14, (2017).

13. Chen, X. et al. In vivo protein turnover rates in varying oxygen tensions nominate MYBBP1A as a mediator of the hyperoxia response. Sci. Adv. 9, eadj4884 (2023).

14. Sørensen, B. S., Busk, M., Overgaard, J., Horsman, M. R. & Alsner, J. Simultaneous Hypoxia and Low Extracellular pH Suppress Overall Metabolic Rate and Protein Synthesis In Vitro. PloS One 10, e0134955 (2015).

15. Rennie, M. J. et al. Muscle protein synthesis measured by stable isotope techniques in man: the effects of feeding and fasting. Clin. Sci. Lond. Engl. 1979 63, 519–523 (1982).

16. Signer, R. A. J., Magee, J. A., Salic, A. & Morrison, S. J. Haematopoietic stem cells require a highly regulated protein synthesis rate. Nature 509, 49–54 (2014).

17. Ja, M. & Raj, S. Developmental Stage-Specific Changes in Protein Synthesis Differentially Sensitize Hematopoietic Stem Cells and Erythroid Progenitors to Impaired Ribosome Biogenesis. Stem Cell Rep. 16, (2021).

18. Huang, Y., Urban, C., Hubel, P., Stukalov, A. & Pichlmair, A. Protein turnover regulation is critical for influenza A virus infection. Cell Syst. 0, (2024).

19. Jovanovic, M. et al. Dynamic profiling of the protein life cycle in response to pathogens. Science 347, 1259038 (2015).

20. Balch, W. E., Morimoto, R. I., Dillin, A. & Kelly, J. W. Adapting proteostasis for disease intervention. Science 319, 916–919 (2008).

21. Hipp, M. S., Kasturi, P. & Hartl, F. U. The proteostasis network and its decline in ageing. Nat. Rev. Mol. Cell Biol. 20, 421–435 (2019).

22. Buchberger, A., Bukau, B. & Sommer, T. Protein quality control in the cytosol and the endoplasmic reticulum: brothers in arms. Mol. Cell 40, 238–252 (2010).

23. Klaips, C. L., Jayaraj, G. G. & Hartl, F. U. Pathways of cellular proteostasis in aging and disease. J. Cell Biol. 217, 51–63 (2018).

24. Pakos-Zebrucka, K. et al. The integrated stress response. EMBO Rep. 17, 1374–1395 (2016).

25. Arimoto, K., Fukuda, H., Imajoh-Ohmi, S., Saito, H. & Takekawa, M. Formation of stress granules inhibits apoptosis by suppressing stress-responsive MAPK pathways. Nat. Cell Biol. 10, 1324–1332 (2008).

26. He, L., Cho, S. & Blenis, J. mTORC1, the maestro of cell metabolism and growth. Genes Dev. 39, 109–131 (2025).

27. Zhao, J., Zhai, B., Gygi, S. P. & Goldberg, A. L. mTOR inhibition activates overall protein degradation by the ubiquitin proteasome system as well as by autophagy. Proc. Natl. Acad. Sci. U. S. A. 112, 15790–15797 (2015).

28. Cui, D. S., Webster, S. M. & Davis, J. H. Integrated proteasomal and lysosomal activity shape mTOR-regulated proteome remodeling. 2024.07.20.603815 Preprint at 10.1101/2024.07.20.603815 (2024).

29. Zhang, Y. et al. Coordinated regulation of protein synthesis and degradation by mTORC1. Nature 513, 440–443 (2014).

30. Neurohr, G. E. et al. Excessive Cell Growth Causes Cytoplasm Dilution And Contributes to Senescence. Cell 176, 1083–1097.e18 (2019).

31. Alber, A. B., Paquet, E. R., Biserni, M., Naef, F. & Suter, D. M. Single Live Cell Monitoring of Protein Turnover Reveals Intercellular Variability and Cell-Cycle Dependence of Degradation Rates. Mol Cell 71, 1079–1091.e9 (2018).

32. Barry, J. D., Donà, Erika, Gilmour, D. & Huber, W. TimerQuant: a modelling approach to tandem fluorescent timer design and data interpretation for measuring protein turnover in embryos. Development 143, 174–179 (2016).

33. Khmelinskii, A. et al. Tandem fluorescent protein timers for in vivo analysis of protein dynamics. Nat. Biotechnol. 30, 708–714 (2012).

34. Alber, A. B. & Suter, D. M. Single-Cell Quantification of Protein Degradation Rates by Time-Lapse Fluorescence Microscopy in Adherent Cell Culture. JoVE J. Vis. Exp. e56604 (2018) doi:10.3791/56604.

35. Bodor, D. L., Rodríguez, M. G., Moreno, N. & Jansen, L. E. T. Analysis of protein turnover by quantitative SNAP-based pulse-chase imaging. Curr. Protoc. Cell Biol. Chapter 8, Unit8.8 (2012).

36. Woodside, K. H. Effects of cycloheximide on protein degradation and gluconeogenesis in the perfused rat liver. Biochim. Biophys. Acta BBA - Gen. Subj. 421, 70–79 (1976).

37. Beugnet, A., Tee, A. R., Taylor, P. M. & Proud, C. G. Regulation of targets of mTOR (mammalian target of rapamycin) signalling by intracellular amino acid availability. Biochem. J. 372, 555–566 (2003).

38. Nandi, D., Woodward, E., Ginsburg, D. B. & Monaco, J. J. Intermediates in the formation of mouse 20S proteasomes: implications for the assembly of precursor beta subunits. EMBO J. 16, 5363–5375 (1997).

39. Khan, S. et al. Immunoproteasomes largely replace constitutive proteasomes during an antiviral and antibacterial immune response in the liver. J. Immunol. Baltim. Md 1950 167, 6859–6868 (2001).

40. Tanaka, K. & Ichihara, A. Half-life of proteasomes (multiprotease complexes) in rat liver. Biochem. Biophys. Res. Commun. 159, 1309–1315 (1989).

41. Cuervo, A. M., Palmer, A., Rivett, A. J. & Knecht, E. Degradation of proteasomes by lysosomes in rat liver. Eur. J. Biochem. 227, 792–800 (1995).

42. Jha, R. K., Kouzine, F. & Levens, D. MYC function and regulation in physiological perspective. Front. Cell Dev. Biol. 11, 1268275 (2023).

43. Huang, M.-J., Cheng, Y., Liu, C.-R., Lin, S. & Liu, H. E. A small-molecule c-Myc inhibitor, 10058-F4, induces cell-cycle arrest, apoptosis, and myeloid differentiation of human acute myeloid leukemia. Exp. Hematol. 34, 1480–1489 (2006).

44. Schwanhäusser, B. et al. Global quantification of mammalian gene expression control. Nature 473, 337–342 (2011).

45. Cerbini, T. et al. Transcription activator-like effector nuclease (TALEN)-mediated CLYBL targeting enables enhanced transgene expression and one-step generation of dual reporter human induced pluripotent stem cell (iPSC) and neural stem cell (NSC) lines. PloS One 10, e0116032 (2015).

46. Ramos-Gonzalez, P. et al. Astrocytic atrophy as a pathological feature of Parkinson’s disease with LRRK2 mutation. Npj Park. Dis. 7, 1–11 (2021).

47. Metzner, C. et al. Superstatistical analysis and modelling of heterogeneous random walks. Nat. Commun. 6, 7516 (2015).

48. Mark, C. et al. Bayesian model selection for complex dynamic systems. Nat. Commun. 9, 1803 (2018).

49. Mark, C., Metzner, C. & Fabry, B. Bayesian inference of time varying parameters in autoregressive processes. Preprint at 10.48550/arXiv.1405.1668 (2014).

50. Martin, B. & Suter, D. M. An out-of-equilibrium definition of protein turnover. BioEssays 45, 2200209 (2023).

51. Padgett, J. & Santos, S. D. M. From clocks to dominoes: lessons on cell cycle remodelling from embryonic stem cells. FEBS Lett. (2020) doi:10.1002/1873-3468.13862.

52. Zatulovskiy, E., Zhang, S., Berenson, D. F., Topacio, B. R. & Skotheim, J. M. Cell growth dilutes the cell cycle inhibitor Rb to trigger cell division. Science 369, 466–471 (2020).

53. Perez-Pinera, P., Ousterout, D. G., Brown, M. T. & Gersbach, C. A. Gene targeting to the ROSA26 locus directed by engineered zinc finger nucleases. Nucleic Acids Res 40, 3741– 3752 (2012).

54. Hara, K. et al. Amino acid sufficiency and mTOR regulate p70 S6 kinase and eIF-4E BP1 through a common effector mechanism. J. Biol. Chem. 273, 14484–14494 (1998).

55. Nielsen, P. J., Manchester, K. L., Towbin, H., Gordon, J. & Thomas, G. The phosphorylation of ribosomal protein S6 in rat tissues following cycloheximide injection, in diabetes, and after denervation of diaphragm. A simple immunological determination of the extent of S6 phosphorylation on protein blots. J. Biol. Chem. 257, 12316–12321 (1982).

56. Price, D. J., Nemenoff, R. A. & Avruch, J. Purification of a hepatic S6 kinase from cycloheximide-treated Rats. J. Biol. Chem. 264, 13825–13833 (1989).

57. Shah, O. J., Anthony, J. C., Kimball, S. R. & Jefferson, L. S. 4E-BP1 and S6K1: translational integration sites for nutritional and hormonal information in muscle. Am. J. Physiol. Endocrinol. Metab. 279, E715–729 (2000).

58. Proud, C. G. mTOR-mediated regulation of translation factors by amino acids. Biochem. Biophys. Res. Commun. 313, 429–436 (2004).

59. Laplante, M. & Sabatini, D. M. Regulation of mTORC1 and its impact on gene expression at a glance. J. Cell Sci. 126, 1713–1719 (2013).

60. Hussein, A. M. et al. Metabolic Control over mTOR-Dependent Diapause-like State. Dev. Cell 52, 236–250.e7 (2020).

61. Bulut-Karslioglu, A. et al. Inhibition of mTor induces a paused pluripotent state. Nature 540, 119–123 (2016).

62. Dörrbaum, A. R., Kochen, L., Langer, J. D. & Schuman, E. M. Local and global influences on protein turnover in neurons and glia. eLife 7, e34202 (2018).

63. Dörrbaum, A. R., Alvarez-Castelao, B., Nassim-Assir, B., Langer, J. D. & Schuman, E. M. Proteome dynamics during homeostatic scaling in cultured neurons. eLife 9, e52939 (2020).

64. Dörrbaum, A. R., Schuman, E. M. & Langer, J. D. Dynamic SILAC to Determine Protein Turnover in Neurons and Glia. in SILAC: Methods and Protocols (ed. Luque-Garcia, J. L.) 1–17 (Springer US, New York, NY, 2023). doi:10.1007/978-1-0716-2863-8_1.

65. Welle, K. A. et al. Time-resolved Analysis of Proteome Dynamics by Tandem Mass Tags and Stable Isotope Labeling in Cell Culture (TMT-SILAC) Hyperplexing. Mol. Cell. Proteomics MCP 15, 3551–3563 (2016).

66. Todorova, P. K. et al. Amino acid intake strategies define pluripotent cell states. Nat. Metab. 6, 127–140 (2024).

67. Whitten, W. K. & Biggers, J. D. Complete development in vitro of the pre-implantation stages of the mouse in a simple chemically defined medium. J. Reprod. Fertil. 17, 399– 401 (1968).

68. Miao, Y., et al. Cycloheximide (CHX) Chase Assay to Examine Protein Half-life. Bio-Protoc. 13, e4690 (2023).

69. Diaz-Cuadros, M., et al. Metabolic Regulation of Species-Specific Developmental Rates. 2021.08.27.457974 https://www.biorxiv.org/content/10.1101/2021.08.27.457974v1 (2021) doi:10.1101/2021.08.27.457974.

70. Matsuda, M. et al. Species-specific segmentation clock periods are due to differential biochemical reaction speeds. Science 369, 1450–1455 (2020).

71. Rayon, T. et al. Species-specific pace of development is associated with differences in protein stability. Science 369, eaba7667 (2020).

72. Suter, D. M. et al. Rapid generation of stable transgenic embryonic stem cell lines using modular lentivectors. Stem Cells Dayt. Ohio 24, 615–623 (2006).

73. Deluz, C. et al. A role for mitotic bookmarking of SOX2 in pluripotency and differentiation. Genes Dev 30, 2538–2550 (2016).

74. Kasparek, P. et al. Efficient gene targeting of the Rosa26 locus in mouse zygotes using TALE nucleases. FEBS Lett. 588, 3982–3988 (2014).

75. Dobin, A. et al. STAR: ultrafast universal RNA-seq aligner. Bioinforma. Oxf. Engl. 29, 15– 21 (2013).

76. Love, M. I., Huber, W. & Anders, S. Moderated estimation of fold change and dispersion for RNA-seq data with DESeq2. Genome Biol. 15, 550 (2014).

77. R Core Team. R: A Language and Environment for Statistical Computing. (R Foundation for Statistical Computing, Vienna, Austria, 2023).

78. Tweedie, S. et al. Genenames.org: the HGNC and VGNC resources in 2021. Nucleic Acids Res. 49, D939–D946 (2021).

79. Rath, S. et al. MitoCarta3.0: an updated mitochondrial proteome now with sub-organelle localization and pathway annotations. Nucleic Acids Res. 49, D1541–D1547 (2021).

80. Wiśniewski, J. R., Zougman, A., Nagaraj, N. & Mann, M. Universal sample preparation method for proteome analysis. Nat. Methods 6, 359–362 (2009).

81. Kulak, N. A., Pichler, G., Paron, I., Nagaraj, N. & Mann, M. Minimal, encapsulated proteomic-sample processing applied to copy-number estimation in eukaryotic cells. Nat. Methods 11, 319–324 (2014).

82. Cox, J. & Mann, M. MaxQuant enables high peptide identification rates, individualized p.p.b.-range mass accuracies and proteome-wide protein quantification. Nat. Biotechnol. 26, 1367–1372 (2008).

83. Cox, J. et al. Accurate proteome-wide label-free quantification by delayed normalization and maximal peptide ratio extraction, termed MaxLFQ. Mol. Cell. Proteomics MCP 13, 2513–2526 (2014).

84. Tyanova, S. et al. The Perseus computational platform for comprehensive analysis of (prote)omics data. Nat. Methods 13, 731–740 (2016).

85. Bendall, S. C. et al. Prevention of amino acid conversion in SILAC experiments with embryonic stem cells. Mol. Cell. Proteomics MCP 7, 1587–1597 (2008).

86. Ershov, D. et al. TrackMate 7: integrating state-of-the-art segmentation algorithms into tracking pipelines. Nat. Methods 19, 829–832 (2022).

87. Weigert, M., Schmidt, U., Haase, R., Sugawara, K. & Myers, G. Star-convex Polyhedra for 3D Object Detection and Segmentation in Microscopy. in 2020 IEEE Winter Conference on Applications of Computer Vision (WACV) 3655–3662 (2020). doi:10.1109/WACV45572.2020.9093435.

88. Munkres, J. Algorithms for the Assignment and Transportation Problems. J. Soc. Ind. Appl. Math. 5, 32–38 (1957).

89. Savitzky, Abraham. & Golay, M. J. E. Smoothing and Differentiation of Data by Simplified Least Squares Procedures. Anal. Chem. 36, 1627–1639 (1964).

90. Pachitariu, M. & Stringer, C. Cellpose 2.0: how to train your own model. Nat. Methods 19, 1634–1641 (2022).

91. Witz, G. guiwitz/microfilm. (2024).

92. Fischler, M. A. & Bolles, R. C. Random sample consensus: a paradigm for model fitting with applications to image analysis and automated cartography. Commun. ACM 24, 381– 395 (1981).

93. Pedregosa, F. et al. Scikit-learn: Machine Learning in Python. J. Mach. Learn. Res. 12, 2825–2830 (2011).

