## Supplementary Text for "Core passive and facultative mTOR-mediated mechanisms coordinate mammalian protein synthesis and decay"

March 3, 2025

### 1 Mammalian Cell-optimized Fluorescent Timer (MCFT)

#### 1.1 Modeling of the MCFT

In the following section, we describe the deterministic model used to quantify the dynamics of the MCFT. The model was already partly developed in our previous work [1]. Briefly, the MCFT consists of the translational fusion of the sfGFP and the mOrange2 fluorescent proteins. Both fluorescent proteins can be in two distinct states, either matured (and thus fluorescent) or not. The two states are coined,"green"($G$)/"red"($R$) (for sfGFP/mOrange2) and "black green" ($B_G$)/"black red"($B_R$) (for sfGFP/mOrange2), respectively. The set of ordinary differential equations (ODEs) describing the time variation in the levels of the different states of the MCFT is given by:

$$\frac{dB_G}{dt} = s - (m_G + k) \times B_G \quad (1)$$

$$\frac{dG}{dt} = m_G \times B_G - k \times G \quad (2)$$

$$\frac{dB_R}{dt} = s - (m_R + k) \times B_R \quad (3)$$

$$\frac{dR}{dt} = m_G \times B_R - k \times R \quad (4)$$

with:

$$k = k_{deg} + k_{dil} \quad (5)$$

$k$ is the decay rate for the MCFT protein. $k_{deg}$ is the protein degradation rate. $k_{dil}$ is the dilution rate, due to cell growth and division. $s$ is the protein synthesis rate. $s$ is an absolute or relative synthesis rate. In the text, we use $S$ when the synthesis rate is normalized to the control condition, such that in the control $S = 1$. $m_G$ and $m_R$ are the maturation rate of the sfGFP and the mOrange2 proteins, respectively. $B_G$, $G$, $B_R$, $R$ are the level of the MCFT in state 'black green", "green", "black red", and "red", respectively. See [1] and Figure 1a-b for more details.

#### 1.2 Steady-state behavior of the MCFT

At steady-state we have:

$$\frac{dB_G}{dt} = 0 \quad (6)$$

$$\frac{dG}{dt} = 0 \quad (7)$$

$$\frac{dB_R}{dt} = 0 \quad (8)$$

$$\frac{dR}{dt} = 0 \quad (9)$$

In other words, we have:

$$B_G^* = \frac{s}{m_G + k} \quad (10)$$

$$G^* = \frac{m_G}{k} \times B_G^* \quad (11)$$

$$B_R^* = \frac{s}{m_R + k} \quad (12)$$

$$R^* = \frac{m_R}{k} \times B_R^* \quad (13)$$

By developing the terms that are linked to observables, we finally reach:

$$G^* = \frac{m_G}{k} \times \frac{s}{m_G + k} \quad (14)$$

$$R^* = \frac{m_R}{k} \times \frac{s}{m_R + k} \quad (15)$$

We can now define the green over red ($G/R$) ratio at steady-state, $\mathcal{R}$:

$$\mathcal{R} = \frac{G^*}{R^*} = \frac{m_G}{m_R} \times \frac{m_R + k}{m_G + k} \quad (16)$$

that only depends on the protein decay rate $k$, $s$ being canceled by the ratio. The decay rate $k$ can thus be computed at steady-state from the $G/R$ ratio, $\mathcal{R}$:

$$k = m_G m_R \times \frac{1 - \mathcal{R}}{m_R \mathcal{R} - m_G} \quad (17)$$

At steady-state, we can then compute the protein synthesis rate $s$ seeing that:

$$s = G^* \times \frac{k(m_G + k)}{m_G} \quad (18)$$

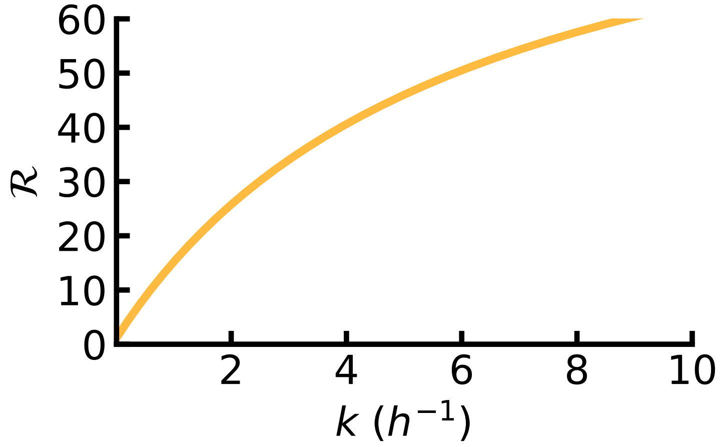

Figure 1: Green over Red ($G/R$) ratio, $\mathcal{R}$, versus the decay rate $k$, at steady-state.

In summary, the measurement of the green and red fluorescences of the MCFT allows to compute the protein synthesis and decay rates, $s$ and $k$, at steady-state. Note that $k$ is in absolute units ($h^{-1}$) while $s$ is in relative units ($fluorescence.h^{-1}$), since it depends on the microscope settings.

##### 1.3 Scaling factor $\alpha$

Since the photon output measured per mature molecule of sfGFP and mOrange2 differ according to microscope settings, we quantified the scaling factor $\alpha$ that allows us to directly compare sfGFP and mOrange2 fluorescence intensities. One way to compute this scaling factor is to induce a transient expression of the MCFT, followed by the removal of the inducer to reach $s = 0$. Doing so and waiting long enough, we can assume that:

$$\frac{dB_G}{dt} = 0 \tag{19}$$

$$\frac{dB_R}{dt} = 0 \tag{20}$$

and:

$$B_G \approx 0 \tag{21}$$

$$B_R \approx 0 \tag{22}$$

This means that when waiting long enough in the absence of newly synthesized proteins, all the MCFT in black-green and black-red states will mature, and thus all MCFT molecules will be fluorescent in both green and red. The dynamics of the green and red states will then follow, under the exact same conditions:

$$\frac{dG}{dt} = -k \times G \tag{23}$$

$$\frac{dR}{dt} = -k \times R \tag{24}$$

We identify two exponential decays:

$$G(t) = G(t=0) \times e^{-kt} \tag{25}$$

$$R(t) = R(t=0) \times e^{-kt} \tag{26}$$

The $G/R$ ratio thus reads:

$$\frac{G(t)}{R(t)} = \frac{G(t=0)}{R(t=0)} \tag{27}$$

And, because the MCFT is a translational fusion of sfGFP and mOrange2 proteins (1:1 ratio in the levels of proteins), we expect:

$$\frac{G(t)}{R(t)} = \frac{G(t=0)}{R(t=0)} = 1 \tag{28}$$

Practically, because fluorescences are given in relative units, this ratio depends on the microscope settings and we have:

$$\frac{G(t)}{R(t)} = \frac{G(t=0)}{R(t=0)} = \alpha \tag{29}$$

This correction factor $\alpha$ is determined experimentally. Importantly, an error on $\alpha$ can propagate on the computation of $s$ and $k$ at steady-state. Indeed, using this correction, the computation of $k$ is done using:

$$k = m_G m_R \times \frac{1 - \frac{\mathcal{R}}{\alpha}}{m_R \frac{\mathcal{R}}{\alpha} - m_G} \tag{30}$$

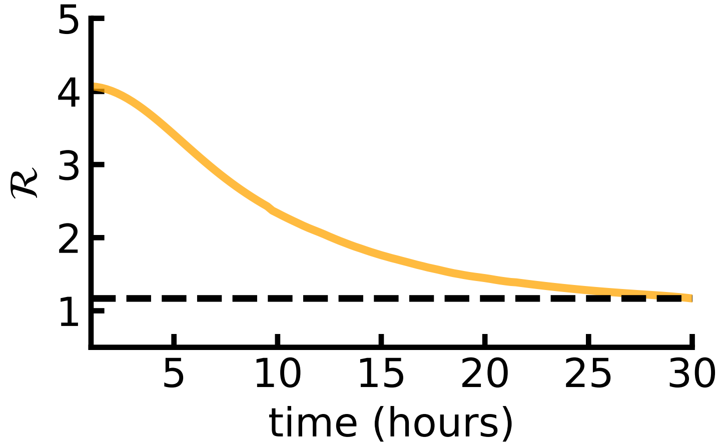

Figure 2: $\mathcal{R}$ decays over time and converges to $\alpha$ when the expression of the MCFT protein is shut down (time t=0h). In this figure $\alpha = 1.2$ and is denoted by the black dotted line.

##### 1.4 Inference of mOrange2 maturation rate $m_R$

To compute $s$ and $k$, we need to know the values of the constants appearing in Figure 1, namely $m_G$, $m_R$, in addition to the already described technical parameter $\alpha$. We assumed that the maturation rate for sfGFP is close to what was reported in the literature ([1, 2] — $m_G \approx 6h^{-1}$). We determined $m_R$ by matching $k$ measured with SNAP chase — in different drug treatment conditions — from the one computed using the G/R ratio. Mathematically, we maximized the log-transformed $\ell_2$-norm:

$$m^*_{R,i} = \arg \max_{m_R \in \Omega_{m_R}} - \ln \left[ (k_{\mathcal{R},i}(m_R) - k_{SNAP,i})^2 \right] \tag{31}$$

for $\Omega_{m_R} = [0.01, 1]$ ($\Omega_{m_R}$ denotes the possible values of $m_R$) discretized in $10^5$ linear bins. $k_{SNAP,i}$ is the degradation rate measured from the SNAP pulse-chase experiment in drug condition $i$. $k_{\mathcal{R},i}(m_R)$ is the degradation rate computed from the timer's green and red fluorescences for a given $m_R$ and condition $i$. $m^*_{R,i}$ is the optimal — i.e. inferred — maturation rate for mOrange2 protein in condition $i$. We then selected the maturation rate value we use, $\hat{m_R}$, as being the median of $m^*_{R,i}$ over conditions $i$.

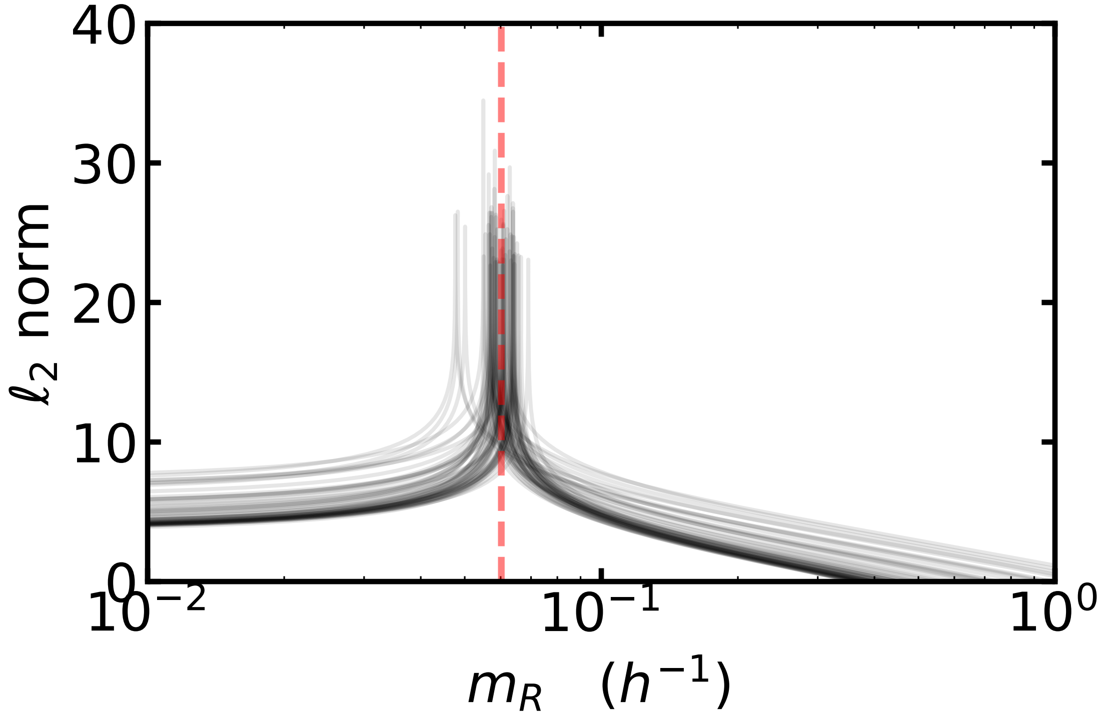

Figure 3: Calibration of the MCFT: determination of the maturation rate of mOrange2 protein, $m_R$. Each gray line represents one condition, different conditions exhibiting different decay rates $k$ used for $m_R^*$ determination. $m_R^*$ is the optimal maturation rate.

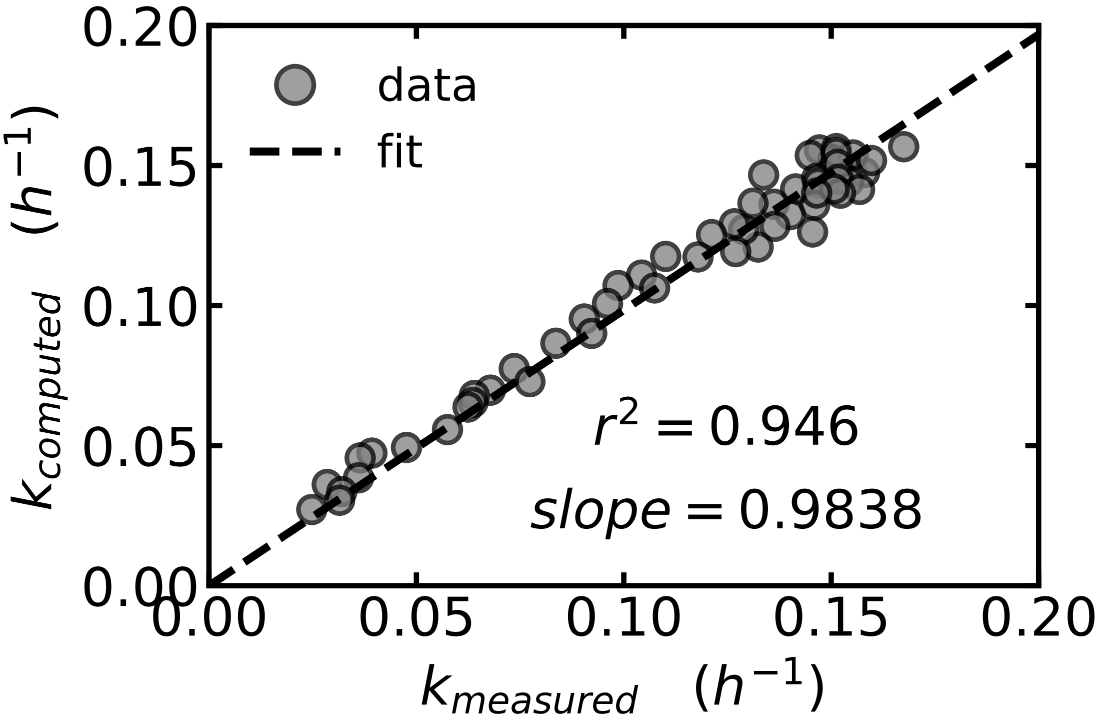

Figure 4: Decay rate computed with $\mathcal{R}$ using $m_R = m_R^*$, $k_{computed}$, versus decay rate measured by SNAP pulse-chase. Dashed black line: best linear fit fixing intercept to 0.

### 2 Hierarchical Bayesian algorithm for inferring the dynamics of time variable rates

#### 2.1 Principle

We adapted a superstatistical Bayesian inference algorithm developed by Fabry et al. [3, 4, 5] for autoregressive models (AR-1) to ODE systems. We computed the (joint) posterior distribution of $s$ and $k$ along with the latent variable $B_G$ (see related section) at each timepoint and propagated this posterior forward and backward along the trajectory. In our case, the trajectories used for the inference are green $G$ and red $R$ fluorescence trajectories acquired by live-cell imaging. Propagation of the posterior distribution relies on two hyperparameters chosen for their ability to recapitulate the data (data retrodiction) and fixed for a whole dataset.

#### 2.2 Likelihood

The posterior distribution is computed from the likelihood. Keeping the same argument as in our original study [1] we choose a likelihood suitable for Gaussian processes — i.e. we assumed normally distributed and independent measurement errors. Therefore, the likelihood $\mathcal{L}$ reads:

$$\mathcal{L} = P(G_o(t), R_o(t)|s(t), k(t), BG_0, t) = \frac{1}{\sigma\sqrt{2\pi}} e^{\frac{-(G_o(t)-G_m(t))^2}{2\sigma^2}} \times \frac{1}{\sigma\sqrt{2\pi}} e^{\frac{-(R_o(t)-R_m(t))^2}{2\sigma^2}} \quad (32)$$

with $\sigma = 20$, which can be considered as a hyperparameter of the algorithm. $G_o(t)$ ($G_m(t)$) and $R_o(t)$ ($R_m(t)$) are the observed (modeled) sfGFP and mOrange2 fluorescent signals at time $t$, respectively.

#### 2.3 Data retrodiction

For data retrodiction, we used inferred time evolution of $s$ and $k$ along with inferred $B_G$ to integrate the ODEs describing the time evolution of timer species levels (see related section). Briefly, we used DifferentialEquations.jl package in Julia to integrate the ODE system using Euler integrator, a time-step of 0.001 hours, and applying forcing functions $s(t)$, $k(t)$, and $B_G(t)$.

#### 2.4 Test on synthetic data

To gain confidence in the ability of our algorithm to infer the time-variation of $s$ and $k$, we generated synthetic timer trajectories with known underlying $s$ and $k$ dynamics. We integrated the set of equations describing timer fluorescence dynamics in-silico using DifferentialEquations.jl in Julia. Tsit5() integrator was used, with a total integration time of 30 hours and a sampling time of 15 min to mimic experimental data. Gaussian noise, $\mathcal{N}(0, 10)$, was then added to the trajectories, to mimic experimental noise. All other model parameters were fixed to biologically plausible values (using for instance those reported in [1]). For all cases tested, including extreme rates time-variations (e.g. dephased sinusoidal variations), we observed a close-to-perfect agreement between inferred time variations and ground truth.

### 3 Passive adaptation model

#### 3.1 General considerations

We assume a protein expression system made of 3 different modules:

- A division machinery, with protein level $A$, protein synthesis rate $s_A$, and protein decay rate $k_A$ — that defines the dilution rate $k_{dil}$;
- A degradation machinery, with protein level $B$, protein synthesis rate $s_B$, and protein decay rate $k_B$;
- A protein of interest (POI), with protein level $C$, protein synthesis rate $s_C$, and protein decay rate $k_C$.

In the following model, we will assume that $A$, $B$, and $C$ dynamics follow:

$$\frac{dA}{dt} = s_A - k_A \times A \tag{33}$$
$$\frac{dB}{dt} = s_B - k_B \times B \tag{34}$$
$$\frac{dC}{dt} = s_C - k_C \times C \tag{35}$$

and that the decay rates for $B$ and $C$, are given by:

$$k_B = k_{deg,B} + k_{dil} \approx k_{dil} \tag{36}$$
$$k_C = k_{deg,C} + k_{dil} = \hat{k}_{deg,C} \times B + k_{dil} \tag{37}$$

Where $\hat{k}_{deg,C}$ is the intrinsic degradation rate of $C$. It means that we assume the proteasome machinery $B$ to be long-lived, decaying only by dilution (see main text for details).

**Importantly**, when $k_{dil} = 0$ (non-dividing and non-infinitively growing cells), $k_B = k_{deg,B}$. Then, we recover the results in the next sections by replacing $k_{dil}$ by $k_{deg,B} = \hat{k}_{deg,B} \times B$ where it has to be replaced.

#### 3.2 Model formalization

Assume that $k_{dil}$ depends on the dynamics (response half-time) of the division machinery ($A$), which ultimately depends on its degradation rate:

$$k_{dil} = k_{deg,A} = \hat{k}_{deg,A} \times B \tag{38}$$

Where $\hat{k}_{deg,A}$ is the intrinsic degradation rate of $A$. According to the previous considerations, we can write that:

$$\frac{dB}{dt} = s_B - \underbrace{\hat{k}_{deg,A} \times B}_{k_{dil}} \times B \tag{39}$$

$$\frac{dC}{dt} = s_C - \underbrace{\hat{k}_{deg,C} \times B}_{k_{deg,C}} \times C - \underbrace{\hat{k}_{deg,A} \times B}_{k_{dil}} \times C \tag{40}$$

Developing the formulas we obtain:

$$\frac{dB}{dt} = s_B - \hat{k}_{deg,A} \times B^2 \tag{41}$$

$$\frac{dC}{dt} = s_C - (\hat{k}_{deg,C} + \hat{k}_{deg,A}) \times B \times C \tag{42}$$

The steady-state solution of the previous set of equations is given by solving $(\dot{B}, \dot{C})|_{(B_{eq},C_{eq})} = \vec{0}$, where $B_{eq}$ and $C_{eq}$ are the steady-state levels of $B$ and $C$ machinery. In other words:

$$0 = s_B - \hat{k}_{deg,A} \times B_{eq}^2 \tag{43}$$

$$0 = s_C - (\hat{k}_{deg,C} + \hat{k}_{deg,A}) \times B_{eq} \times C_{eq} \tag{44}$$

That implies (keeping only positive solutions):

$$B_{eq} = \left( \frac{s_B}{\hat{k}_{deg,A}} \right)^{\frac{1}{2}} \tag{45}$$

$$C_{eq} = \frac{s_C}{\hat{k}_{deg,C} + \hat{k}_{deg,A}} \times \frac{1}{B_{eq}} \tag{46}$$

Meaning that:

$$C_{eq} = \frac{s_C}{\hat{k}_{deg,C} + \hat{k}_{deg,A}} \times \left( \frac{s_B}{\hat{k}_{deg,A}} \right)^{-\frac{1}{2}} \tag{47}$$

Now if we assume that all the synthesis rates are proportional to a so-called global synthesis rate $s$:

$$s_A \propto s_B \propto s_C \propto s$$

And assume that the global synthesis rate $s$ is changed — perturbed — by a scaling factor $\beta$:

$$\hat{s} = s \times \beta$$

Where $\hat{s}$ is the perturbed $s$. This finally implies that:

$$\hat{s}_A \propto \hat{s}_B \propto \hat{s}_C \propto \hat{s}$$

.

Or written differently:

$$\hat{s} = s \times \beta \tag{48}$$
$$\hat{s}_A = s_A \times \beta \tag{49}$$
$$\hat{s}_B = s_B \times \beta \tag{50}$$
$$\hat{s}_C = s_C \times \beta \tag{51}$$

Interestingly, we thus have:

$$C_{eq}|_{\hat{s}} = s_C \times \beta \times \frac{1}{\hat{k}_{deg,C} + \hat{k}_{deg,A}} \times \left( \frac{\hat{k}_{deg,A}}{\beta \times s_B} \right)^{\frac{1}{2}} = \beta \times C_{eq}|_s \tag{52}$$

And importantly:

$$C_{eq}|_{\hat{s}} = \frac{\hat{s}_C}{\hat{k}_{deg,C} + \hat{k}_{deg,A}} \times \frac{\sqrt{\beta}}{B_{eq}|_s} \tag{53}$$

and defining the effective decay rate $\tilde{k}_C$ we obtain:

$$C_{eq}|_{\hat{s}} = \frac{\hat{s}_C}{\tilde{k}_C} \tag{54}$$

with:

$$\tilde{k}_C = (\hat{k}_{deg,C} + \hat{k}_{deg,A}) \times \frac{B_{eq}|_s}{\sqrt{\beta}} = \tilde{k}_C|_s \times \frac{1}{\sqrt{\beta}} \tag{55}$$

which finally implies:

$$\frac{k_B|_s}{\tilde{k}_C} = \sqrt{\beta} \tag{56}$$

Finally, the equilibrium protein level $C_{eq}$ fold-change is given by:

$$\frac{C_{eq}|_{\hat{s}}}{C_{eq}|_s} = \sqrt{\beta} \tag{57}$$

In the main text, we denoted by $S$ the fold-change in $s$ with respect to the control condition. Thus $S = \frac{s}{s_{control}} = \beta$.

##### 3.3 In-silico passive adaptation model simulation

To validate our analytical developments, we integrated the set of equations describing passive adaptation until reaching steady state. The integration was performed using DifferentialEquations.jl in Julia. Tsit5() integrator was used. We observed (Figure 5) a perfect correspondence between simulation results and analytical predictions for $k_f/k_i \equiv k_B|_s/\tilde{k}_B$ and $P_f/P_i \equiv B_{eq}|_{\hat{s}}/B_{eq}|_s$.

##### 3.4 Modelling the adaptation of protein decay to changes in protein synthesis

###### 3.4.1 No adaptation model

In this model, we assume that only the protein synthesis rate $s$ is changing upon CHX treatment. Using the same notations as previously, $s$ is changing by a factor $\beta$. At equilibrium, the protein level is thus changing as:

$$\frac{[P]_f}{[P]_i} = \frac{\beta \times s_i}{k} \times \frac{k}{s_i} = \beta \tag{58}$$

Where $[P]_f$, $[P]_i$ is the final, respectively initial, protein concentration. Similarly, $s_f$, $s_i$ is the final, respectively initial, protein synthesis rate. In this case, the protein decay rate $k$ is constant.

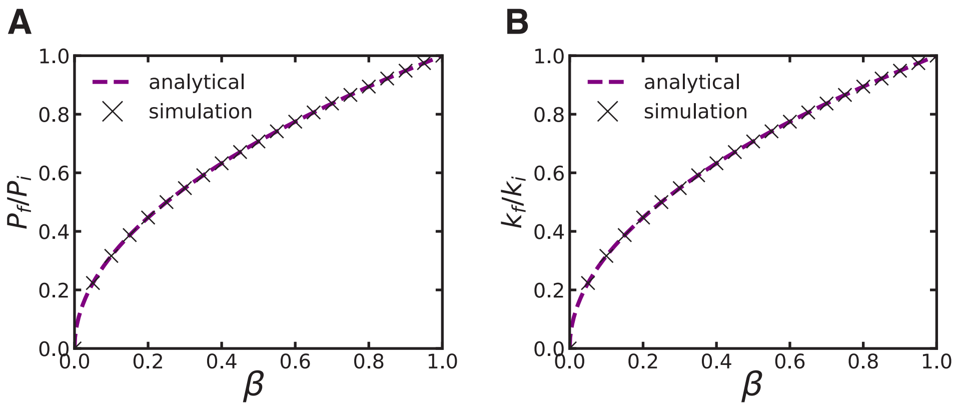

Figure 5: Passive adaption model predictions of the fold-changes in protein level and decay rate. **A.** Predicted fold-change in the protein level $P_f/P_i$ with respect to the fold-change in synthesis rate $\beta$. **B.** Predicted fold-change in the decay rate $k_f/k_i$ with respect to the fold-change in synthesis rate $\beta$.

##### 3.4.2 Perfect adaptation model

In this model, we assume that the decay rate $k$ adapts to $s$ in a manner that allows to fully maintain protein levels. In this case, we will have, trivially:

$$\frac{[P]_f}{[P]_i} = 1 \tag{59}$$

and, subsequently:

$$FI(k) = \frac{k_f}{k_i} = \beta \tag{60}$$

#### 3.5 Change in proteome content in the passive adaptation model: absolute abundance, relative abundance, and concentration

Here we describe how the passive adaptation model should theoretically affect proteome composition in terms of protein concentrations, absolute and relative abundances. As shown previously,

$$[P_i]_{t_f} = \sqrt{\beta} \times [P_i]_{t_i} \tag{61}$$

where $[P_i]_{t_f}$ and $[P_i]_{t_i}$ are the protein concentrations for protein $i$ at the final and initial time, respectively. From this relation, we can derive changes in absolute protein abundance:

$$\frac{P_{i,t_f}}{V_f} = \sqrt{\beta} \times \frac{P_{i,t_i}}{V_i} \tag{62}$$

where $V_f$ and $V_i$ are the cell volume at the final and initial time, respectively. Rearranging the equation we have:

$$P_{i,t_f} = \sqrt{\beta} \times \frac{V_f}{V_i} \times P_{i,t_i} \tag{63}$$

Now, we define the relative abundance of protein $i$, $\sigma_i$ as:

$$\sigma_i = \frac{P_i}{\sum_i P_i} \tag{64}$$

We can compute this relative abundance at time $t_i$ and $t_f$:

$$\begin{aligned}
\sigma_{i,t_i} &= \frac{P_{i,t_i}}{\sum_i P_{i,t_i}} && (65)\\
\sigma_{i,t_f} &= \frac{P_{i,t_f}}{\sum_i P_{i,t_f}} && (66)\\
&= \frac{\sqrt{\beta} \times \frac{V_f}{V_i} \times P_{i,t_i}}{\sum_i \sqrt{\beta} \times \frac{V_f}{V_i} \times P_{i,t_i}} && (67)\\
&= \frac{P_{i,t_i}}{\sum_i P_{i,t_i}} && (68)\\
&= \sigma_{i,t_i} && (69)
\end{aligned}$$

Finally, if both $k_{deg}$ and $k_{dil}$ perfectly follow the passive adaptation model, their changes will scale linearly upon changes in $s$. As a consequence, the relative abundance of all proteins stays constant.

#### 3.6 Including the ubiquitination pathway into the passive adaptation model

##### 3.6.1 The ubiquitination pathway

Here we introduce a new species $U$ standing for the ubiquitination machinery in the passive adaptation model. In this model, protein degradation depends on two subsequent steps:

protein ubiquitination and proteasomal degradation of the ubiquitinated protein. The set of ODEs describing the time-evolution of this system reads as:

$$\frac{dA}{dt} = s_A - k_A \times A^2 \tag{70}$$

$$\frac{dP}{dt} = s - k_U \times U \times P \tag{71}$$

$$\frac{dU}{dt} = s_U - k_U \times U \times P - k_{d,U} \times A \times U \tag{72}$$

$$\frac{d[UP]}{dt} = k_U \times U \times P - k_{UP} \times A \times [UP] \tag{73}$$

Applying the steady-state condition:

$$\left.\frac{dA}{dt}\right|_{A^*} = 0 \tag{74}$$

$$\left.\frac{dP}{dt}\right|_{P^*} = 0 \tag{75}$$

$$\left.\frac{dU}{dt}\right|_{Uj} = 0 \tag{76}$$

$$\left.\frac{d[UP]}{dt}\right|_{[UP]^*} = 0 \tag{77}$$

We obtain the following steady-state state:

$$A^* = \sqrt{\frac{s_A}{k_A}} \tag{78}$$

$$P^* = \frac{s}{k_U} \times \frac{k_{d,U \times A^*}}{s_u - s} \tag{79}$$

$$U^* = \frac{s_U - s}{k_{d,U} \times A^*} \tag{80}$$

$$UP^* = \frac{s}{k_U A^*} \tag{81}$$

with the restriction $(s_U - s) > 0$. Scaling all the synthesis rates by $\beta$ we obtain:

$$A^*|_\beta = \sqrt{\beta} \times A^* \tag{82}$$

$$P^*|_\beta = \sqrt{\beta} \times P^* \tag{83}$$

$$U^*|_\beta = \sqrt{\beta} \times U^* \tag{84}$$

$$UP^*|_\beta = \sqrt{\beta} \times UP^* \tag{85}$$

The total protein content $P_{tot}$ is, in this case

$$P_{tot} = P + UP \tag{86}$$

since the two species are undistinguishable in the experiments. Thus, it follows that:

$$P^*_{tot}|_\beta = \sqrt{\beta} \times P^*_{tot} \tag{87}$$

In other words, we recover the results for the passive adaptation model that does not take into account the ubiquitination step in the protein degradation pathway.

### 4 Mass spectrometry data analysis

#### 4.1 Half-life and $psk_{deg}$ of NIH/3T3 proteins

The $psk_{deg}$ was computed using the dynamic SILAC we performed in NIH/3T3 (see after).

#### 4.2 Predicting changes in protein relative amount for $psk_{deg}$

##### 4.2.1 Tuning the passive adaptation model

To predict the changes in protein relative amount for $psk_{deg}$ (hereafter we drop the $ps$ for simplicity), taking into account the observed offset between the adaptation of $k_{deg}$ and $k_{dil}$, we can rewrite the equation of the passive adaptation model taking $k_{dil}$ as an input:

$$k_{dil} \equiv input$$

That is, the dilution rate is given by the observation. In this case, we can rewrite the previous differential equations, keeping the same symbols:

$$\frac{dB}{dt} = s_B - k_{dil}B \tag{88}$$

$$\frac{dC}{dt} = s_C - \hat{k}_{deg,C} \times B \times C - k_{dil} \times C \tag{89}$$

At steady-state, $(\dot{B}, \dot{C}) = \vec{0}$ and:

$$B^* = \frac{s_B}{k_{dil}} \tag{90}$$

$$C^* = \frac{s_C}{\hat{k}_{deg,C} \times B^* + k_{dil}} \tag{91}$$

where $B^*$ and $C^*$ are the steady-state protein level for the degradation machinery and the POI, respectively.

Now assume the global synthesis rate $s$ is changed by a factor $\beta$ between an initial (subscript $i$) and a final (subscript $f$) state. We have:

$$\frac{B_f^*}{B_i^*} = \beta \quad (92)$$

$$\frac{C_f^*}{C_i^*} = \beta \frac{\hat{k}_{deg,C} \times B_i^* + k_{dil,i}}{\hat{k}_{deg,C} \times \beta B_i^* + k_{dil,f}} \quad (93)$$

and, by definition, we have:

$$\frac{k_f}{k_i} = \frac{\beta k_{deg,C,i} + k_{dil,i}}{k_{deg,C,i} + k_{dil,f}} \quad (94)$$

###### 4.2.2 The short-lived protein limit

When $k_{dil} \approx 0$ ($k_{dil} \ll k_{deg,C}$) we obtain the following limit case:

$$\frac{C_f^*}{C_i^*} = 1 \quad (95)$$

$$\frac{k_f}{k_i} = \beta \quad (96)$$

###### 4.2.3 The long-lived protein limit

When $k_{deg,C} \approx 0$ ($k_{dil} \gg k_{deg,C}$) we obtain the following limit case:

$$\frac{C_f^*}{C_i^*} = \beta \quad (97)$$

$$\frac{k_f}{k_i} = 1 \quad (98)$$

###### 4.2.4 Change in proteome content in the tuned passive adaptation model: absolute abundance, relative abundance, and concentration

In this section, we will show that, taking into account the offset in adaptation between $k_{deg}$ and $k_{dil}$ observed in NIH/3T3:

- the fold-change in protein level will be dependent on the protein-specific degradation rate ;

- the relative abundance of a given protein (its stoichiometry) will depend upon its protein-specific degradation rate.

As stated previously, the relative abundance of a given protein $i$, $\sigma_i$, reads:

$$\sigma_i = \frac{P_i}{\sum_i P_i} \tag{99}$$

We can compute this relative abundance at time $t_i$ and $t_f$:

$$\begin{aligned}
\sigma_{i,t_i} &= \frac{P_{i,t_i}}{\sum_i P_{i,t_i}} && (100)\\
\sigma_{i,t_f} &= \frac{P_{i,t_f}}{\sum_i P_{i,t_f}} && (101)\\
&= \beta P_{i,t_i} \frac{\frac{k_{deg,i}\times + k_{dil,i}}{\beta\times k_{deg,i}+k_{dil,f}}}{\sum_i P_{i,t_f}} && (102)\\
& && (103)
\end{aligned}$$

### 5 Dynamic SILAC analysis

#### 5.1 Recycling probability of light amino acids

We performed `search B` in MaxQuant as described in [7] with modifications given in STAR Methods. From the `evidence.txt` output of MaxQuant we first filtered out all the peptides assigned to contaminants or decoy sequences, i.e. we removed all peptides for which `Reverse` and `Potential contaminant` column are equal to `+`. We then extracted peptides containing two Arg/Lys residues using the condition `Missed cleavage == 1`. We determined the number of peptides that contain two heavy Arg/Lys (HH) or one heavy and one light Arg/Lys (HL) by counting the number of peptides matching, respectively, the condition `Modifications == "Arg10 as ptm, Lys8 as ptm"` or `Modifications == "Arg10 as ptm" or "Lys8 as ptm"`. Taking into account other possible modifications (Acetyl, Oxidation) did not change the results dramatically. We then computed the probability of heavy amino acid incorporation $P(H)$:

$$P(H) = \frac{2 \times \frac{HH}{HL}}{1 + 2 \times \frac{HH}{HL}} \tag{104}$$

with $HH$ or $HL$ the number of peptides with two heavy Arg/Lys or one light and one heavy Arg/Lys, respectively.

##### 5.2 Determination of protein decay rate

We performed `search A` in MaxQuant as described in [7], with modifications given in STAR Methods. From the `peptides.txt` output of MaxQuant we first filtered out all the peptides assigned to contaminants or decoy sequences, i.e. we removed all peptides for which `Reverse` and `Potential contaminant` column are equal to `+`. We then removed all peptides showing false heavy signal at timepoint 0 hours: e.g., we removed all peptides for which `Intensity H DMSO_t0` $> 0$. From this, we computed the fraction of remaining light peptides $\%L$:

$$\%L = \frac{1}{1 + \frac{H}{L}} \tag{105}$$

with $\frac{H}{L}$ that is given by, e.g., the `Ratio H/L DMSO_t2` column. We computed the fraction of remaining old peptides, $\%old$, taking into account the recycling probability of light amino acids $P(H)$. Using the following equation:

$$\%old = 1 - \frac{1 - \%L}{P(H)^{MC+1}} \tag{106}$$

where $MC$ is the number of missed cleavage site for a given peptide. Finally, as described in Material and Methods, we log-transformed the $\%old$ data and fitted a linear model with respect to time, such that: $ln(\%old) = -k \times t$ (canonical exponential decay).

#### References

[1] Alber, A. B., Paquet, E. R., Biserni, M., Naef, F. & Suter, D. M. Single Live Cell Monitoring of Protein Turnover Reveals Intercellular Variability and Cell-Cycle Dependence of Degradation Rates. *Mol Cell* **71**, 1079–1091.e9 (2018).

[2] Balleza, E., Kim, J. M. & Cluzel, P. Systematic characterization of maturation time of fluorescent proteins in living cells. *Nature methods* **15**, 47–51 (2018). URL `https://www.ncbi.nlm.nih.gov/pmc/articles/PMC5765880/`.

[3] Mark, C. *et al.* Bayesian model selection for complex dynamic systems. *Nature Communications* **9**, 1803 (2018). URL `https://www.nature.com/articles/s41467-018-04241-5`. Number: 1 Publisher: Nature Publishing Group.

[4] Metzner, C. *et al.* Superstatistical analysis and modelling of heterogeneous random walks. *Nature Communications* **6**, 7516 (2015).

[5] Mark, C., Metzner, C. & Fabry, B. Bayesian inference of time varying parameters in autoregressive processes (2014). URL http://arxiv.org/abs/1405.1668. ArXiv:1405.1668 [q-bio].

[6] Schwanhäusser, B. *et al.* Global quantification of mammalian gene expression control. *Nature* **473**, 337–342 (2011). URL https://www.nature.com/articles/nature10098. Number: 7347 Publisher: Nature Publishing Group.

[7] Dörrbaum, A. R., Schuman, E. M. & Langer, J. D. Dynamic SILAC to Determine Protein Turnover in Neurons and Glia. In Luque-Garcia, J. L. (ed.) *SILAC: Methods and Protocols*, 1–17 (Springer US, New York, NY, 2023). URL https://doi.org/10.1007/978-1-0716-2863-8_1.
