## Supplementary Figures and Tables for "Core passive and facultative mTOR-mediated mechanisms coordinate mammalian protein synthesis and decay"

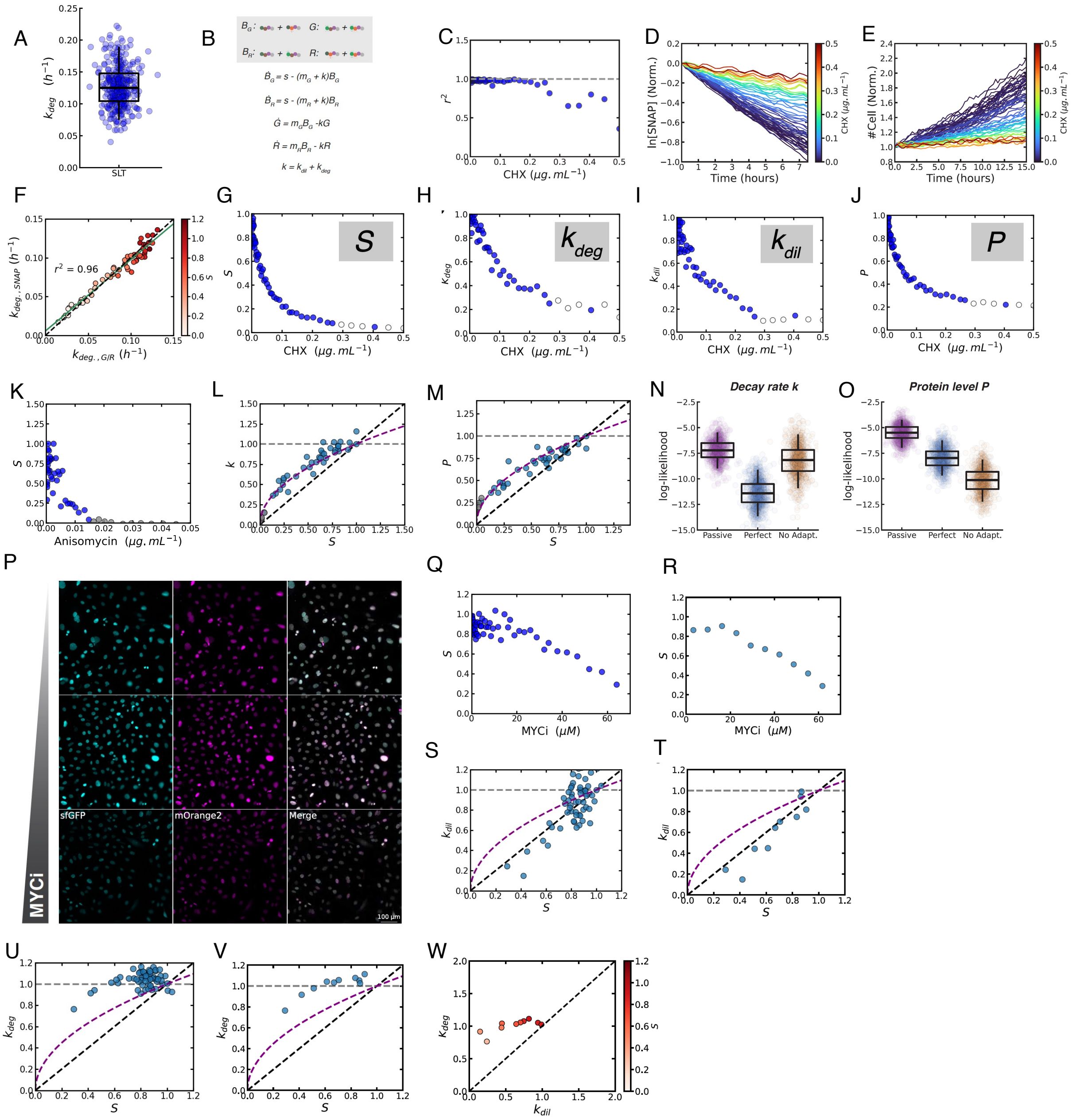


**Figure S1:** (A) Single cell $k_{deg}$ of the SLT measured by SNAP pulse-chase labeling. Boxes: interquartile range; horizontal line: median; vertical lines: 5-95 percentiles. N = 324 lineages (single cells or mother & computationally-fused daughter cells, see Methods). (B) Set of ordinary differential equations describing changes in green and red fluorescence of the MCFT depending on the protein synthesis (*S*) and decay (*k*) rates. $m_G$ and $m_R$: maturation rates for sfGFP and mOrange2, respectively. $B_G$: Black green, i.e. sfGFP protein that did not mature yet; $B_R$: Black red; i.e. mOrange2 protein that did not mature yet. *G*: Green (mature sfGFP); *R*: Red (mature mOrange2). (C) Coefficient of determination $r^2$ of the fit between cell number trajectories and exponential division model for the different CHX concentrations used. (D) SNAP pulse-chase labeling after 48 h of treatment with different CHX concentrations. Traces are the normalized (to t = 0 h), log-transformed, sum of the intensities of all segmented nuclei per condition. (E) Normalized (to t = 0 h) cell number over time after 48 h of CHX treatment. Color scale: CHX concentration for each condition. (F) $k_{deg}$ computed using SNAP pulse-chase trajectories ($k_{deg,SNAP}$) vs using G/R ratios of the MCFT ($k_{deg,G/R}$) for different values of *S*. Values of *S* were normalized to *S* without CHX treatment. x=y black dashed line: perfect concordance between the two ways of computing $k_{deg}$. Green plain line: best linear fit ($r^2$ = 0.96) between $k_{deg,SNAP}$ and $k_{deg,G/R}$. Color bar: fold-change in S for different CHX concentrations. (G-J) Fold-change in *S* (G), $k_{deg}$ (H), $k_{dil}$ (I), and *P* of the SLT (J), with respect to CHX concentration. K, Fold-change in synthesis rate *S* as a function of anisomycin concentration. (L) Fold-change in *k* as a function of *S* upon anisomycin treatment. (M) Fold-change in *P* as a function of *S* upon anisomycin treatment. K-M, Gray dots: data points for which exponential division was lost. (N-O) Log-likelihood for the different models, using both decay rate *k* (N) and protein level *P* (O) data. (P) Snapshots in green (cyan) and red (magenta) fluorescence for NIH/3T3 cell populations treated with different MYCi concentrations (from top to bottom: 1, 14.4, and 64 µM) for 48 hours. (Q-R) Fold-change in *S* with respect to MYCi concentration, unbinned (Q) and binned (R) data. (S-V), Fold-change in $k_{dil}$ (S,T) and $k_{deg}$ (U,V) versus fold-change in *S* upon MYCi treatment, unbinned (S,U) and binned (T,V) data. W, Fold-change in $k_{deg}$ with respect to the fold-change in $k_{dil}$ upon MYCi treatment. Black dashed lines: prediction for perfect adaptation; Gray dashed lines: prediction for no adaptation. Purple dashed curved lines: prediction for passive adaptation. (W) Color bar: fold-change in S for different MYCi concentrations. x=y diagonal black dashed line: equal fold-change in degradation and dilution rates. The values shown for *S*, *P*, $k_{deg,}$ and $k_{dil}$ are normalized on the respective values for control conditions.

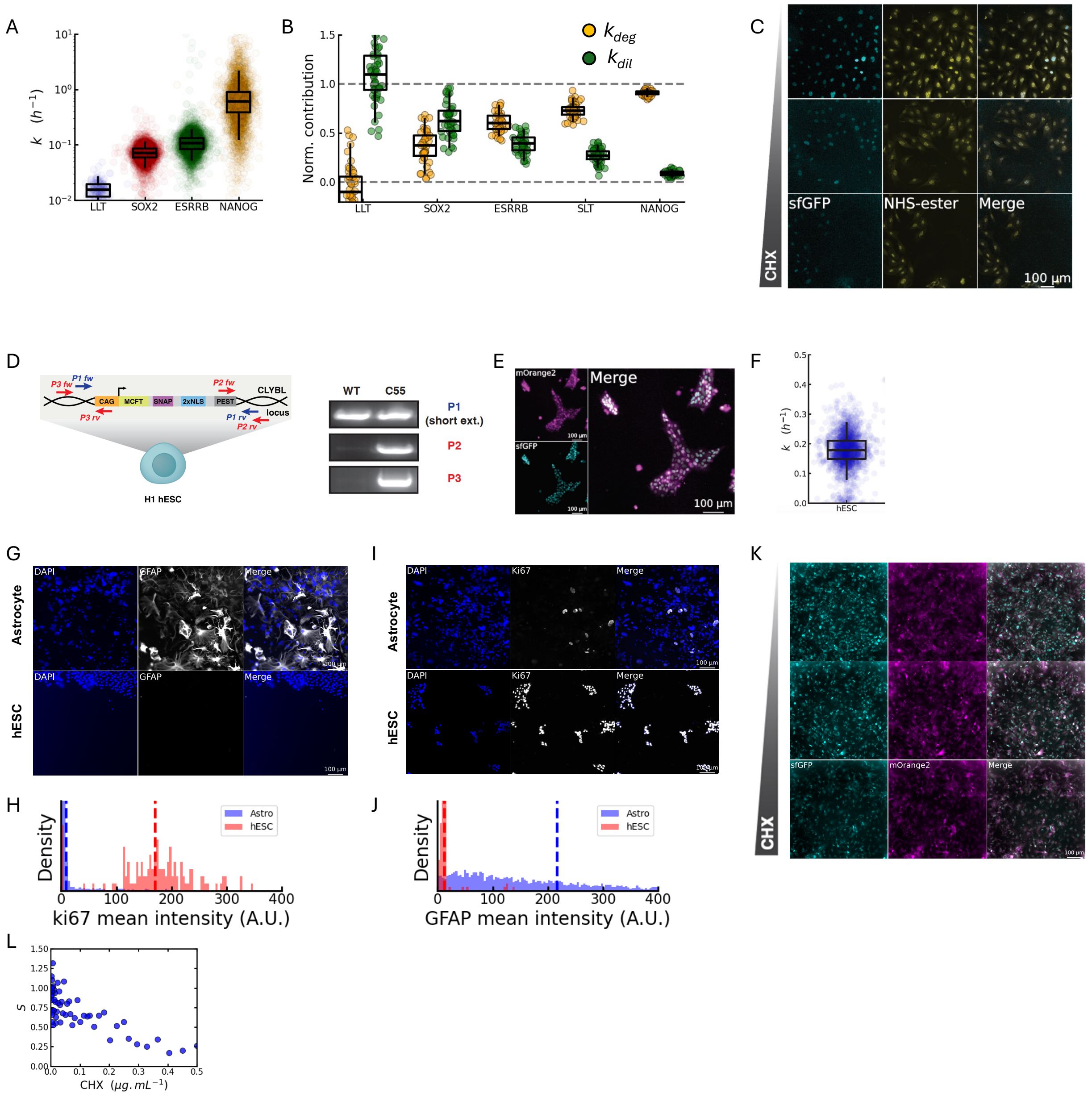


**Figure S2: Characterization of NIH/3T3, hES cell lines, and hESC-derived astrocyte-enriched cultures.** (A) Single cell decay rates of the LLT (blue), SOX2 (red), ESRRB (green), and NANOG (orange) proteins. Boxes: interquartile range; horizontal line: median; vertical line: 5-95 percentiles. N = circa 150 (LLT), 3500 cells otherwise. (B) Normalized contributions of $k_{deg}$ (yellow) and $k_{dil}$ (green) to $k$ for a given $S$ (see Methods). Boxes: interquartile range; horizontal line: median; vertical line: 5-95 percentiles. Horizontal gray dashed lines: theoretical minimum (y=0) and maximum (y=1) possible values. (C) Snapshots in the green (sfGFP - magenta) and NHS-ester channels (yellow) of NIH/3T3 SLT cell populations treated with different CHX concentrations (from top to bottom: 0.008, 0.1125, and 0.5 µg/mL) for 6 days. (D) Left: Scheme of the MCFT construct knocked into H1 hESCs at the CLYBL locus. Right: PCR genotyping of the knock-in clones. P1, P2, and P3 refer to the set of primers used for PCR. A band in the upper panel indicates the amplification of at least one WT locus. WT: wild-type; C55: selected clone knocked-in into the CLYBL locus with the SLT. (E) Snapshot of the hESC line harboring the SLT (sfGFP: cyan; mOrange2: magenta). (F) $k$ of the SLT measured using G/R ratio in the hESC knock-in cell line. Boxes: interquartile range; horizontal line: median; vertical line: 5-95 percentiles. N = 2142 cells. (G-J)

Immunofluorescence for Ki67 (G) and GFAP (I) proteins. Densities of Ki67 (H) and GFAP (J) immunofluorescence signal mean intensities in astrocyte-enriched (blue) and hESC (red) cultures. The vertical dashed lines represent the mean of the related density. (K) Green (cyan) and red (magenta) fluorescence snapshots of astrocyte-enriched cultures treated with different CHX concentrations (from top to bottom: 0.008, 0.1125, and 0.5 μg/mL) for 48 hours. (L) Fold-change in synthesis rate $S$ as a function of CHX concentration in astrocyte-enriched cultures.

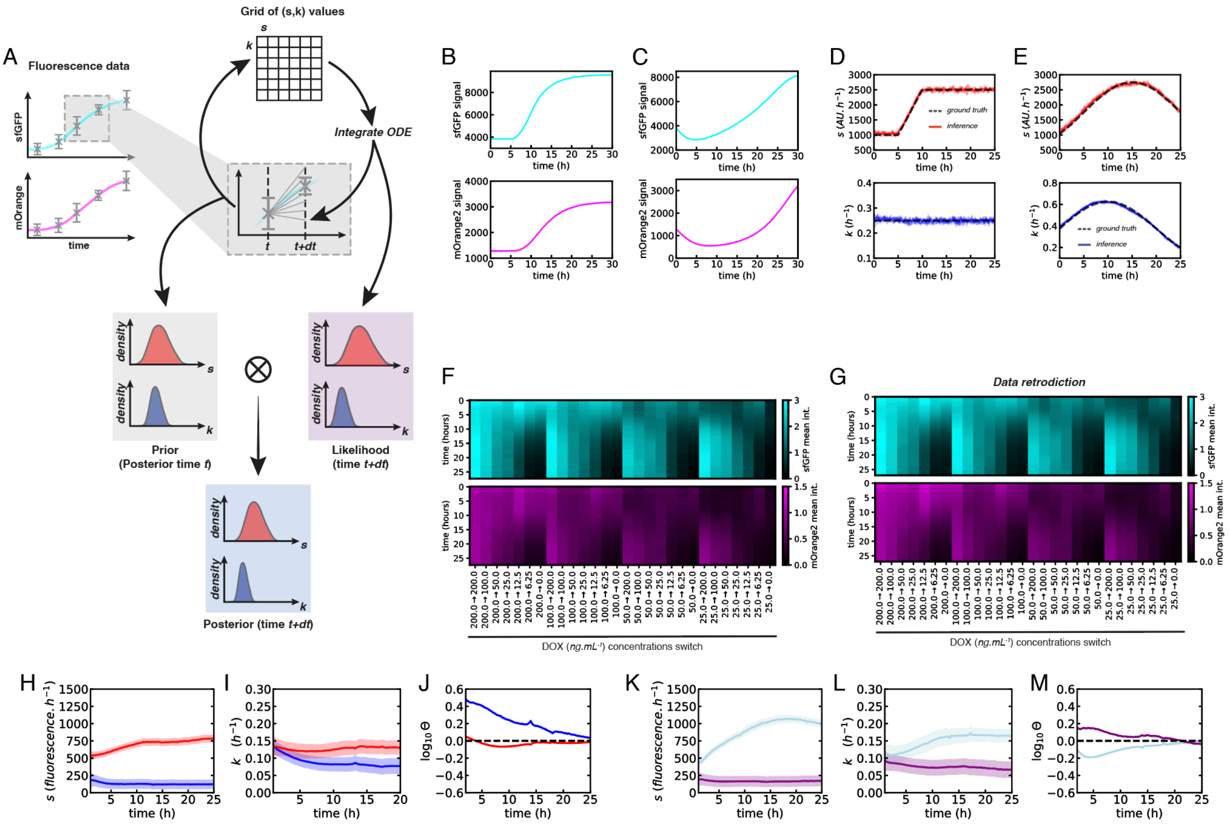

**Figure S3: Determining temporal variations in *S* and *k* using a hierarchical superstatistical Bayesian inference algorithm.** (A) Scheme of the inference strategy. sfGFP (cyan) and mOrange2 (magenta) fluorescence trajectories are both used in the inference. The algorithm iteratively uses each point at time *t* and *t+dt* of the fluorescence trajectory to compute a likelihood for the values of *S* and *k* at time *t*. The convolution of the likelihood distribution with the prior distribution (i.e. the posterior distribution computed at time *t*) allows obtaining the posterior distribution for *S* and *k* for time *t+dt*. This process is then iterated on timepoints *t+dt* and *t+2dt*, etc., for all data points available. (B) sfGFP (cyan) and mOrange2 (magenta) fluorescence trajectories generated in-silico for a linear increase of *S* between 5 and 10 hours. (C) sfGFP (cyan) and mOrange2 (magenta) fluorescence trajectories generated in silico for phase-shifted sinusoidal variations of *S* and *k*. (D) Inferences of the time variation of *S* and *k* from the trajectories in (B). (E) Inferences of the time variation of *S* and *k* from the trajectories in (C). (F) Smoothed sfGFP (cyan) and mOrange2 (magenta) trajectories for experiments with changes in dox concentrations. (G) Data retrodiction for sfGFP (cyan) and mOrange2 (magenta) fluorescence trajectories. (H-M), *S* (H,K), *k* (I,L), and imbalance (J,M) trajectories inferred from MCFT measurements during MYCi pulse (H-J) or release (K-M). Red: control conditions. Dark blue: MYCi pulse. Purple: MYCi treatment. Light blue: MYCi release. (H-I) and (K-L), plain lines: average of the posterior distribution; shaded regions: SD of the posterior distribution.

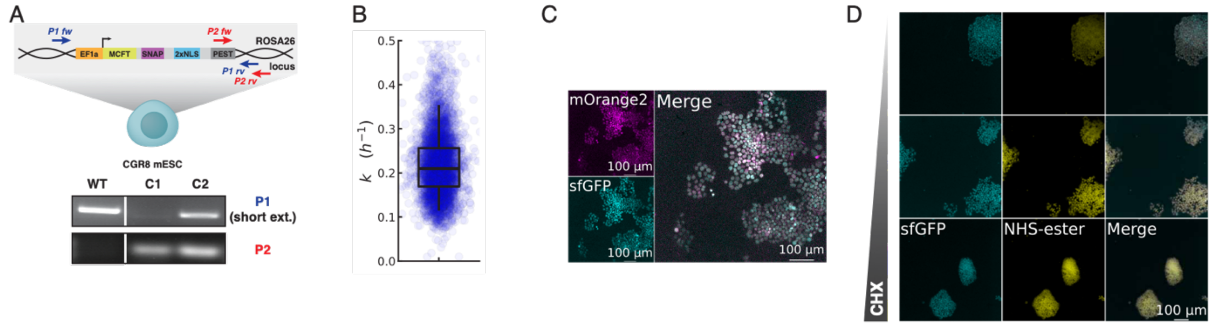


**Figure S4: Characterization of the mESC SLT knock-in cell line.** (A) Scheme of the SLT construct knocked into CGR8 cells at the ROSA26 locus. 2xNLS: two copies of a Nuclear Localization Signal. PCR genotyping of the knock-in clones. P1 and P2 refer to the set of primers used for PCR. A band in the upper panel indicates the amplification of at least one WT locus. WT: wild-type CGR8 mESCs; C1, C2: clones knocked into the ROSA26 locus with the SLT. An irrelevant lane was sliced from the gel picture. C1 was selected for all experiments. (B) *k* of the SLT measured using G/R ratio in the mESC knock-in cell line. Boxes: interquartile range; horizontal line: median; vertical line: 5-95 percentiles. N = 4790 cells. (C) Snapshot of green (cyan) and red (magenta) fluorescence of the mESC line harboring the SLT. (D) Snapshots in the sfGFP (cyan) and NHS-ester (yellow) channels of mESC SLT cell populations treated with different CHX concentrations (from top to bottom: 0.008, 0.1125, and 0.5 µg/mL) for 48 hours.

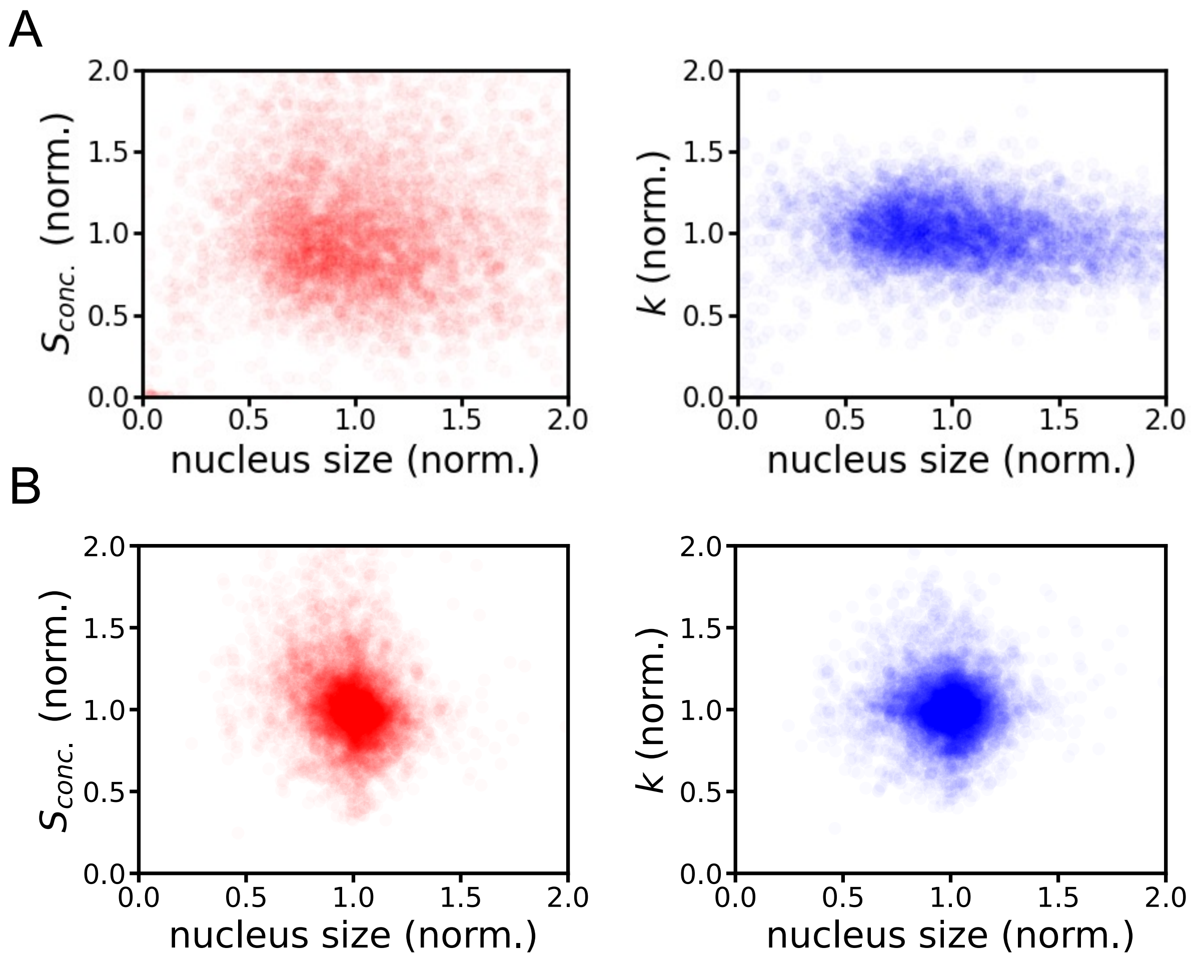


**Figure S5: $S_{conc.}$ and $k$ do not correlate with nucleus size.** All MCFT NIH/3T3 cell lines presented in the main text were used (A). For mESC, the MCFT-Pspc1 cell line was used (B). N > 7000 cells.

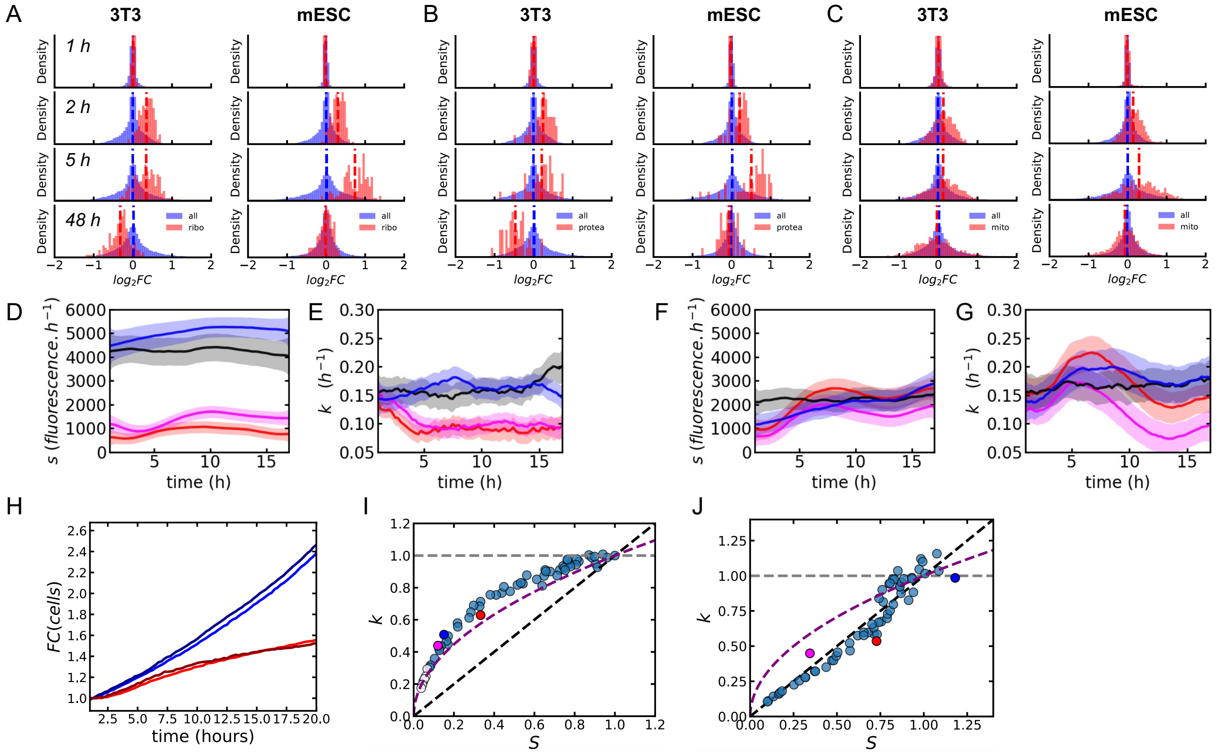

**Figure S6:** (A-C) Density of fold-changes in RNA levels for ribosomal (A), proteasomal (B), and mitochondrial (C) transcripts, with respect to all other transcripts. The vertical dashed lines represent the mean of the related density. (D-G), *S* (D,F) and *k* (E,G) trajectories inferred from MCFT measurements during CHX pulse in NIH/3T3 (D-E) and mES (F-G) cells in the presence or absence of 200 nM ISRIB. Black: control conditions. Blue: ISRIB treatment. Red: CHX treatment. Pink: CHX + ISRIB treatment. The average (plain line) and the SD (shaded region) of the posterior distribution are represented for each timepoint. (H) Normalized (to t = 0 h) cell number over time during CHX pulse in mESC in the presence or absence of INK128. Blue: control condition. Dark blue: INK128 treatment. Red: CHX treatment. Brown: CHX + ISRIB treatment. (I-J) Fold-change in *k* as a function of *S* upon CHX treatment (17 hours – the endpoint of the CHX pulse experiment) in the presence or absence of INK128 in NIH/3T3 (I) and mES (J) cells. Light blue: datapoints from CHX plate experiment (see Figures 1 and 4). Dark blue: INK128 treatment. Red: CHX treatment. Pink: CHX + INK128 treatment.

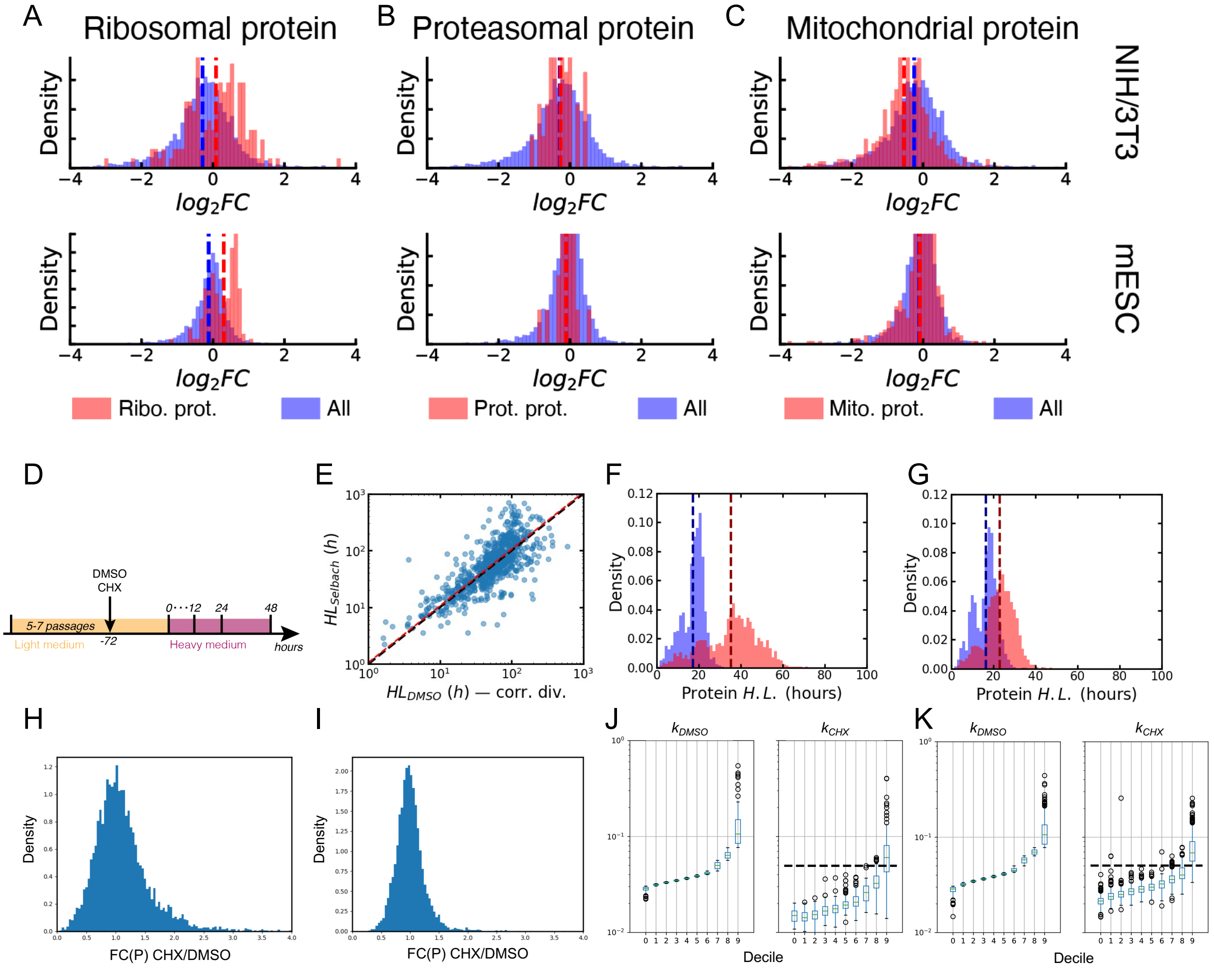


**Figure S7: Dynamic SILAC for NIH/3T3 and mES cells treated with 0.05 µg/mL CHX.** (A-C) Density of fold-changes in protein levels for ribosomal (A), proteasomal (B), and mitochondrial (C) proteins, with respect to all other proteins. The vertical dashed lines represent the mean of the related density. (D) Scheme of the dynamic SILAC experiment (timepoints 1, 2, 4, and 6 hours are not shown). (E) Correlation between published and computed protein half-life in NIH/3T3 cells. (F-G) Density of protein half-lives for unique proteins shared between DMSO (blue) and CHX (red) conditions in NIH/3T3 (F) and mES (G) cells. (H-I), Distribution of protein level fold-changes between CHX and DMSO conditions quantified by LFQ on the first timepoint of the dSILAC in NIH/3T3 (H) and mES (I) cells. (J-K) Measured decay rate in DMSO and CHX conditions, for NIH/3T3 (J) and mESC (K), binned according to ps$k_{deg}$ in DMSO condition. The horizontal black dotted line represents the average decay rate in the DMSO condition.

### Supplementary Tables

| | | | | | | |
|---|---|---|---|---|---|---|
| 0.5 | 0.45 | 0.405 | 0.3645 | 0.32805 | 0.295245 | 0.2657205 |
| 0.25 | 0.225 | 0.2025 | 0.18225 | 0.164025 | 0.1476225 | 0.13286025 |
| 0.125 | 0.1125 | 0.10125 | 0.091125 | 0.0820125 | 0.07381125 | 0.06643013 |
| 0.0625 | 0.05625 | 0.050625 | 0.0455625 | 0.04100625 | 0.03690563 | 0.03321506 |
| 0.03125 | 0.028125 | 0.0253125 | 0.02278125 | 0.02050313 | 0.01845281 | 0.01660753 |
| 0.015625 | 0.0140625 | 0.01265625 | 0.01139063 | 0.01025156 | 0.00922641 | 0.00830377 |
| 0.0078125 | 0.00703125 | 0.00632813 | 0.00569531 | 0.00512578 | 0.0046132 | 0.00415188 |
| 0.00390625 | 0.00351563 | 0.00316406 | 0.00284766 | 0.00256289 | 0.0023066 | 0.00207594 |

**Supplementary Table 1: CHX concentration used for the steady-state experiment.** Concentrations are given in µg/mL, following the layout used to perform the experiment in 96-well plates. The highlighted CHX concentrations are the ones used to show representative microscopy snapshots in the main figures.

| | | | | | | |
|---|---|---|---|---|---|---|
| 0.05 | 0.045 | 0.0405 | 0.03645 | 0.032805 | 0.0295245 | 0.02657205 |
| 0.025 | 0.0225 | 0.02025 | 0.018225 | 0.0164025 | 0.01476225 | 0.01328603 |
| 0.0125 | 0.01125 | 0.010125 | 0.0091125 | 0.00820125 | 0.00738113 | 0.00664301 |
| 0.00625 | 0.005625 | 0.0050625 | 0.00455625 | 0.00410063 | 0.00369056 | 0.00332151 |
| 0.003125 | 0.0028125 | 0.00253125 | 0.00227813 | 0.00205031 | 0.00184528 | 0.00166075 |
| 0.0015625 | 0.00140625 | 0.00126563 | 0.00113906 | 0.00102516 | 0.00092264 | 0.00083038 |
| 0.00078125 | 0.00070313 | 0.00063281 | 0.00056953 | 0.00051258 | 0.00046132 | 0.00041519 |
| 0.00039063 | 0.00035156 | 0.00031641 | 0.00028477 | 0.00025629 | 0.00023066 | 0.00020759 |

**Supplementary Table 2: Anisomycin concentration used for the steady-state experiment.** Concentrations are given in µg/mL, following the layout used to perform the experiment in 96-well plates.

| | | | | | | |
|---|---|---|---|---|---|---|
| 64 | 57.6 | 51.84 | 46.656 | 41.9904 | 37.79136 | 34.012224 |
| 32 | 28.8 | 25.92 | 23.328 | 20.9952 | 18.89568 | 17.006112 |
| 16 | 14.4 | 12.96 | 11.664 | 10.4976 | 9.44784 | 8.503056 |
| 8 | 7.2 | 6.48 | 5.832 | 5.2488 | 4.72392 | 4.251528 |
| 4 | 3.6 | 3.24 | 2.916 | 2.6244 | 2.36196 | 2.125764 |
| 2 | 1.8 | 1.62 | 1.458 | 1.3122 | 1.18098 | 1.062882 |
| 1 | 0.9 | 0.81 | 0.729 | 0.6561 | 0.59049 | 0.531441 |
| 0.5 | 0.45 | 0.405 | 0.3645 | 0.32805 | 0.295245 | 0.2657205 |

**Supplementary Table 3: MYCi concentration used for the steady-state experiment.** Concentrations are given in µM, following the layout used to perform the experiment in 96-well plates. The highlighted MYCi concentrations are the ones used to show representative microscopy snapshots in the main figures.

| name | sequence (5'->3') | reference |
|---|---|---|
| CLYBL wild type genotyping primer, forward | TGACTAAACACTGTGCCCCA | Fernandopulle et al. 2018 |
| CLYBL wild type genotyping primer, reverse | AGGCAGGATGAATTGGTGGA | Fernandopulle et al. 2018 |
| CLYBL 5’ insert genotyping primer, forward | CAGACAAGTCAGTAGGGCCA | Fernandopulle et al. 2018 |
| CLYBL 5’ insert genotyping primer, reverse | AGAAGACTTCCTCTGCCCTC | Fernandopulle et al. 2018 |
| CLYBL 3’ insert genotyping primer, forward | CCGCTCTTTGGAGAAGGTAA | This study |
| CLYBL 3’ insert genotyping primer, reverse | GAACGATTTACTGGGCAGTC | Nickolls et al. 2020 |
| ROSA26 3' insert genotyping primer, forward | AATACCTTTCTGGGAGTTCT | This study |
| ROSA26 3' insert genotyping primer, forward | GCACTTGCTCTCCCAAAGTC | Kasparek et al. 2014 |
| ROSA26 3' wild type genotyping primer, reverse | GGCGGATCACAAGCAATAAT | Kasparek et al. 2014 |

**Supplementary Table 4: List of primers used for genotyping the cell lines generated in this study.**

### Supplementary Movies

**Supplementary Movie 1: CHX (0.1 µg/mL) pulse for NIH/3T3 SLT.**

**Supplementary Movie 2: CHX (0.1 µg/mL) pulse release NIH/3T3 SLT.**

**Supplementary Movie 3: CHX (0.1 µg/mL) pulse for hESC SLT.**

**Supplementary Movie 4: CHX (0.1 µg/mL) release for hESC SLT.**

**Supplementary Movie 5: CHX (0.1 µg/mL) pulse for mESC SLT.**

**Supplementary Movie 6: CHX (0.1 µg/mL) release for mESC SLT.**

For all Supplementary movies, one field of view is shown. One snapshot was taken every 15 minutes in the green and red fluorescence channels. Images are flat field-corrected and background-subtracted (Methods).
